# Multiple adaptive trajectories shape antimalarial resistance evolution in Ethiopian *Plasmodium falciparum*

**DOI:** 10.64898/2026.11.09.764565

**Authors:** Leen N. Vanheer, Metmiku Yohannes, Dagnachew Fenta Workie, Amanuel Shimelash, Meklit Shiferaw, Dawit Hailu Alemayehu, Yisehak Teshome, Shewafera Ademu, Nina Billows, Fekadesilassie Mozart Mesafint, Fikregabrail Aberra Kassa, Melat Abdo, Yisihak Abreham Weldehana, Natnael Lemessa Hundera, Haimanot Andualem Ayalsew, Migbaru Keffale Bezabih, Daniel Neafsey, Bryan Greenhouse, Susana Campino, Nicholas Hathaway, Fitsum Girma Tadesse

## Abstract

*Plasmodium falciparum* malaria continues to impose a substantial public health burden across sub-Saharan Africa, with Ethiopia experiencing a recent resurgence in malaria cases. We analysed whole genome data from 120 *P. falciparum* isolates collected in 2024 across five ecologically and epidemiologically diverse transmission settings in Ethiopia and placed these in the context of 1,220 publicly available genomes. Parasite populations were highly structured within Ethiopia, while Horn of Africa isolates formed a genetically distinct regional cluster relative to East and West African populations. We identified evidence of recent positive selection surrounding the artemisinin resistance associated *Pf*K13 R622I variant and detected co-occurrence with *Pfcrt* mutations together with recurrent selection signals at genes within the *Pf*K13-associated haemoglobin endocytosis pathway, including *Pfmca2*, *Pfkic7*, *Pfkic8* and *Pfkic9*. We further observed the emergence of the *Pf*PX1 D384A-PIN haplotype, with shared identity-by-descent linking Ethiopian and Ugandan carriers, supporting a recent introduction followed by local selection. Widespread *Pfhrp2/3* deletions occurred on diverse genetic backgrounds. Together, these findings reveal a complex adaptive landscape shaped by antimalarial and diagnostic selection pressures and suggest that parasite evolution in the Horn of Africa proceeds through multiple adaptive trajectories, rather than through the spread of a single dominant lineage.

## Introduction

*Plasmodium falciparum* remains a leading cause of morbidity and mortality across sub-Saharan Africa, imposing a substantial public health burden [1]. After major gains up until 2015, progress has since stalled and, in some settings, reversed. Ethiopia has experienced a resurgence of malaria [2], largely driven by *P. falciparum*, eroding years of local progress. This rebound has occurred in the context of multiple interacting pressures, including population displacement and disruption of health services associated with conflict, environmental and climate changes [3], insecticide resistance in malaria vectors [4], and the spread of invasive mosquito species. At the same time, emerging parasite adaptations threaten effectiveness of key malaria control tools. These include the emergence of artemisinin partial resistance (ART-R) and the spread of *P. falciparum* parasites evading rapid diagnostic tests (RDT) through histidine-rich protein 2 and 3 (*Pfhrp2/3*) gene deletions.

Unlike chloroquine and sulfadoxine-pyrimethamine resistance that spread to Africa from Southeast Asia [7,8], ART-R has arisen *de novo* at multiple African sites, particularly in East Africa and the Horn of Africa [9]. ART-R is driven mainly by single nucleotide polymorphisms (SNPs) in the *Pfkelch13 (Pfk13)* gene beta-propeller region, which impair haemoglobin endocytosis [10–12], leading to reduced parasite susceptibility and delayed parasite clearance after artemisinin-based combination therapy (ACT) [12]. Several ART-R associated *Pfk13* alleles have emerged locally and expanded regionally: including R622I, first observed in northwest Ethiopia in 2013–2014 [13], later in Eritrea [14], and parts of Sudan [15]; R561H, first reported in Rwanda in 2014 [10] and now clonally expanding across neighbouring countries in the Great Lakes region [15–17]; C469Y, A675V, and P441L, which were initially described in Uganda in 2016–2017 [18] and have since shown varying degrees of regional expansion, with P441L now reported in multiple African countries [18,19]. This mosaic pattern of geographically distinct resistance associated alleles suggests multiple independent origins of ART-R in Africa and highlights the need not only to track the emergence, spread, and evolutionary dynamics of *Pf*K13 variants, but also to understand the broader genomic contexts that facilitate their establishment and expansion.

Artemisinin resistance is increasingly recognized as a complex trait influenced by variation at multiple loci beyond *Pfk13*. Additional loci involved in *Pf*K13-mediated haemoglobin endocytosis pathways, including Kelch13-interacting proteins *Pf*KIC1-12, *Pf*UBP1, *Pf*MCA2, *Pf*EPS15 and *Pf*AP-2μ [12,20,21], *Pf*Coronin and *Pf*Falcipain-2a, have been associated with reduced artemisinin susceptibility [22–24], suggesting that resistance may be shaped by a broader genomic background.

Evidence of delayed *in vivo* parasite clearance linked to *Pfk13* mutations is now well established across the region. Although artemether-lumefantrine (AL) remains effective across Africa, recent reports of declining efficacy raise concerns about partner drug resistance [25]. A recent study implicated *Pfpx1* with reduced lumefantrine and dihydroartemisinin susceptibility in Uganda [26], underscoring the need for molecular surveillance beyond *Pfk13*. Concurrently, the convergence of *Pfk13* R622I and *Pfhrp2/3* deletions in the Horn of Africa poses a dual threat to malaria control by compromising both treatment efficacy and parasite detection [27,28]. Co-endemic *Plasmodium vivax* further complicates the malaria landscape in the region: chloroquine and primaquine use for radical cure can sustain selection pressure on *P. falciparum* through residual drug exposure [29]. Together, these forces may produce a distinct resistance profile in the region. Previous genomic studies have revealed marked genetic diversity and regional differentiation among *P. falciparum* isolates in Ethiopia, reflecting complex transmission dynamics [30–32]. However, publicly available whole genomes from Ethiopia remain few [11,33], constraining fine-scale inference of population structure, genome-wide diversity and emerging selective processes that could inform surveillance. Comprehensive, time-resolved genomic surveillance is therefore needed to characterize the genetic backgrounds in which resistance and diagnostic-evasion mutations arise and spread across the region.

To address this gap, we generated whole genome sequence (WGS) data from 120 *P. falciparum* isolates collected in 2024 from five ecologically and epidemiologically diverse districts in Ethiopia. These data were used to characterize population structure, detect signatures of selection, and examine genomic background associated with resistance-linked variants and *Pfhrp2/3* deletions.

## Results

### Dataset generation and genomic variant discovery

To characterize the genomic landscape of *P. falciparum* in Ethiopia within a broader regional context, we generated whole genome data for 123 Ethiopian isolates collected in 2024 across five sites (**Figure 1A**), derived from two studies (**Supplementary Figure S1**). After excluding three isolates with high levels of missingness, 120 newly sequenced Ethiopian genomes were retained for downstream analyses. These genomes were analysed alongside publicly available whole genome data. After removal of 134 isolates due to missingness, the comparative publicly available dataset comprised 1340 high-quality genomes from Ethiopia (n=161), Eritrea (n=175), Sudan (n=67), Ghana (n=100), Kenya (n=199), Mali (n=78), Nigeria (n=66), Tanzania (n=211), Uganda (n=163), India (n=98), Myanmar (n=2), and Thailand (n=20) (**Table S1**, **Supplementary Data 1**). Sequencing quality metrics, including percentage of reads mapped and coverage across chromosomes, are provided in **Supplementary Figure S2**. Across the African isolates, 1,106,484 high-quality SNPs were identified, of which 294,224 were bi-allelic variants with a within-country minor allele frequency ≥ 0.01, providing dense resolution for population genetic analysis.

**Figure 1.**
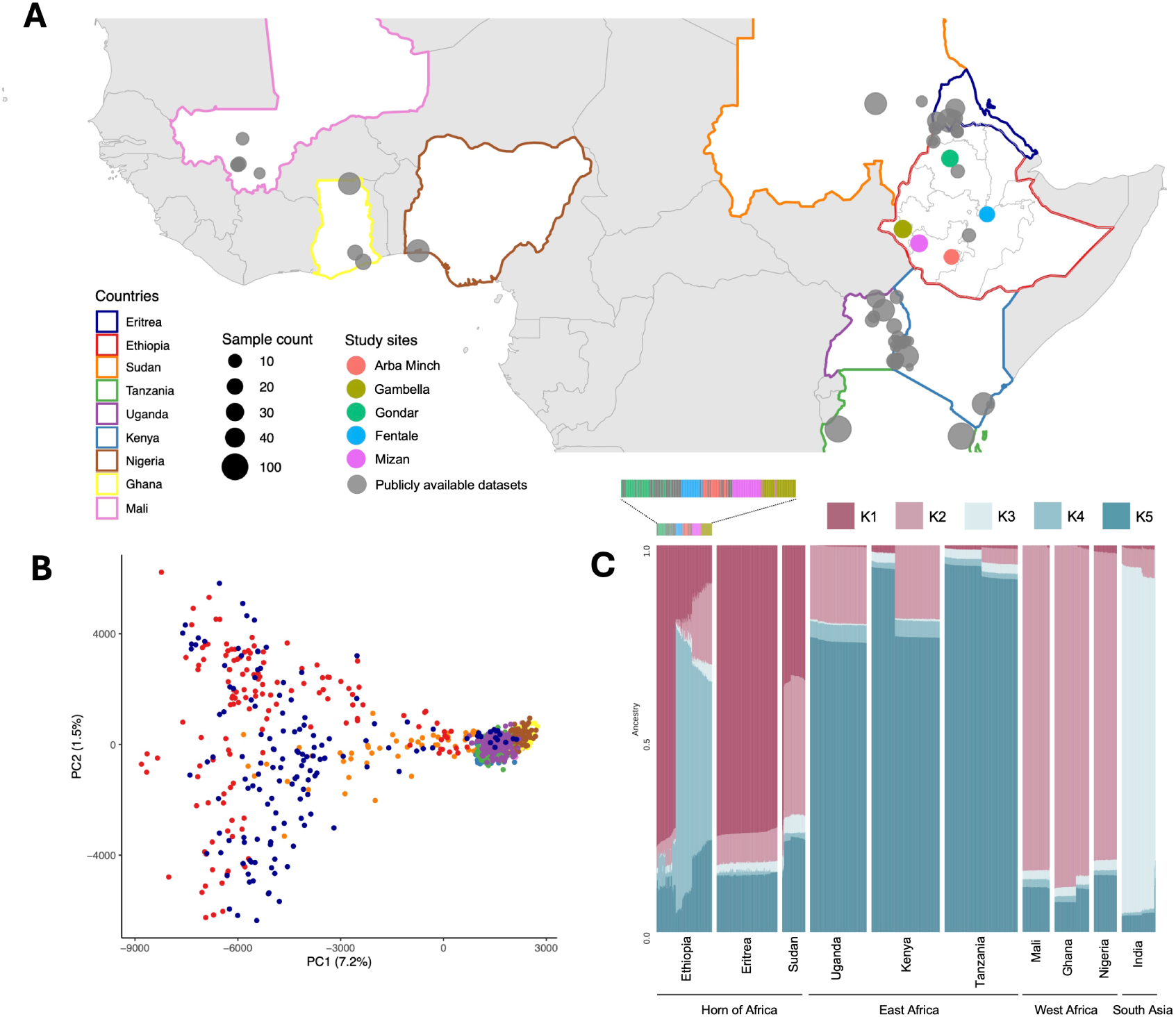
Ethiopian *P. falciparum* isolates in the context of African population structure. Population structure and ancestry composition of 1318 *Plasmodium falciparum* isolates, including 120 Ethiopian isolates sequenced in this study and 1198 publicly available genomes. (**A**) Map of African countries and sites from which genomes were included in this analysis. Country names are indicated by the country outline colour, circle size represents sample count, and fill colour denotes the study site for the newly sequenced Ethiopian isolates. (**B**) SNP-based PCA of African isolates, coloured by country. For polyclonal samples, the dominant allele was selected. Identical or very similar isolates were removed (of a cluster with IBD > 0.95, only one sample was kept). (**C**) Admixture ancestral proportion per isolate (column) for each country, coloured by ancestral populations (K). Analysis for K= 2-10 is presented in **Supplementary Figure S4**. Colour bar above Ethiopian isolates indicates collection sites, as presented in (A).

### Horn of Africa isolates form a genetically distinct cluster

To situate newly sequenced Ethiopian isolates within the broader regional genomic landscape, we compared them against 1100 publicly available *P. falciparum* genomes from the Horn of Africa, East Africa and West Africa (**Figure 1A, Table S1**). SNP-based PCA showed that isolates from the Horn of Africa, including Ethiopia, Eritrea and Sudan, form a genetically distinct cluster compared to East and West African populations (**Figure 1B).**

Notably, when Horn of Africa isolates were excluded, East and West African genomes resolved into more clearly separated clusters **(Supplementary Figure S3**), further underscoring the genetic distinctiveness of the Horn of Africa population compared to other African regions. To assess broader regional relationships beyond Africa, 98 Indian isolates were included in the analysis. These formed a distinct cluster separate from the Horn of Africa parasites (**Supplementary Figure S3**), showing substantial genetic differentiation between the two populations. Consistent with these clustering patterns, genome-wide admixture analysis further highlighted the unique ancestry composition of Horn of Africa isolates compared with those from East and West Africa and India. Within Ethiopia, isolates from the Northern region displayed ancestry profiles closely aligned with Eritrean and Sudanese isolates, whereas isolates from the remaining Ethiopian sites displayed a largely distinct ancestral composition not widely observed elsewhere in the sampled regions of Africa (**Figure 1C, Supplementary Figure S4**). Together, these findings indicate that Horn of Africa *P. falciparum* populations occupy a genetically differentiated position within the broader African parasite landscape, with evidence of both cross-border connectivity in the north and locally distinctive ancestry components within Ethiopia.

### High genetic heterogeneity among Ethiopian *P. falciparum* isolates

Building on this regional context, we next resolved the within-country population structure of Ethiopian isolates. Ethiopian parasite populations exhibited substantial genetic heterogeneity, with marked variation in the proportion of polyclonal infections across sites. F_ws_ based estimates showed the lowest proportion of polyclonal infections in Gondar (23.1%), whereas the remaining sites showed higher levels ranging from 35.7% in Mizan to 47.4% in Fentale (**Figure 2A-B**), consistent with spatial heterogeneity in transmission intensity. PCA revealed strong geographic structuring among Ethiopian parasite populations, broadly recapitulating the spatial relationships between the sites (**Figure 2C**).

**Figure 2.**
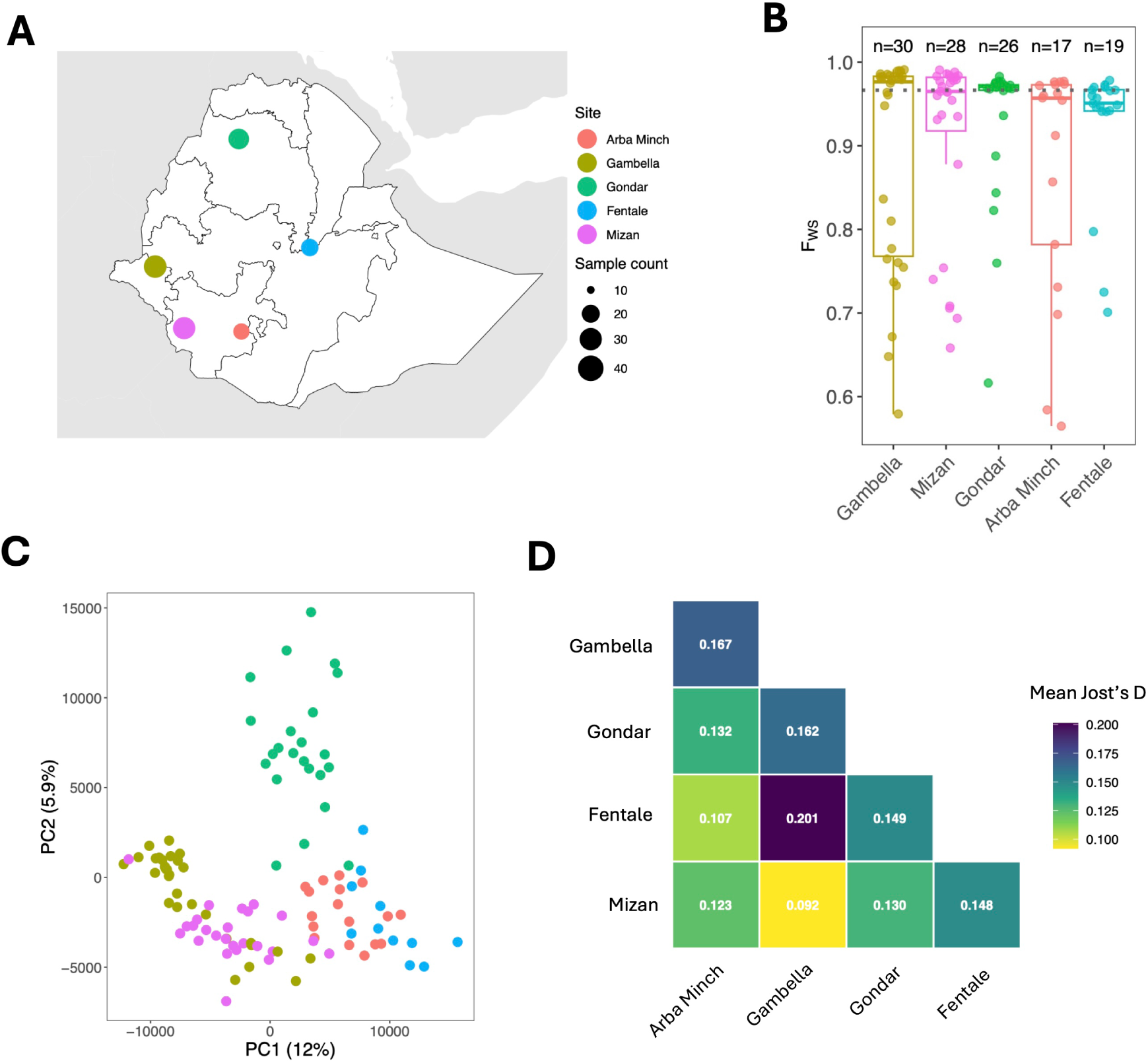
Population structure and differentiation of 120 Ethiopian isolates collected and sequenced in this study. **(A)** Map of Ethiopia indicating the five study sites. The sample count is presented by the size of the circle at each site. (**B**) Within-host diversity estimated by the in-breeding coefficient (F_ws_) per study site. (**C**) Population structure was examined by SNP-based principal component analysis (PCA) coloured by study site. The dominant allele was selected for polyclonal samples. Identical or very similar isolates were removed (of a cluster with IBD > 0.95, only one sample was kept). (**D**) Population differentiation was explored by calculating Jost’s D in comparisons between collection sites, using the multi-allelic VCF. Heatmap showing mean Jost’s D across high heterozygosity markers in comparisons between collection sites. Positions with high Jost’s D can be found in **Supplementary Data 2.**

Population differentiation analysis using Jost’s D analysis identified 183 SNPs exhibiting high genetic differentiation (Jost’s D > 0.7) across pairwise comparisons between Ethiopian collection sites (**Supplementary Data 2**). The greatest divergence was observed between Fentale and Gambella (**Figure 2D**), highlighting pronounced ecological and epidemiological contrasts within Ethiopia. Notably, several highly differentiated SNPs were located within the *Pfcrt* gene, a key determinant of chloroquine resistance, particularly in comparisons involving Gambella versus Arba Minch and Gambella versus Fentale (**Supplementary Data 2)**, suggestive of site-specific selective pressures linked to historical drug use.

### Distribution of molecular markers of drug resistance and *Pfhrp2/3* deletions

Two WHO validated *Pf*K13 beta-propeller variants were detected among Ethiopian isolates: R622I, present in 50% of Gondar isolates (13/26), and A675V, identified in two isolates from Gambella (**Figure 3A, Supplementary Figure S5**) [34]. Beyond emerging markers of ART-R, substantial geographic heterogeneity was observed at other drug resistance loci, particularly *Pfcrt*, a key determinant of chloroquine resistance. Geographic differences in *Pf*CRT K76T frequencies broadly reflected patterns of *P. vivax* co-endemicity and chloroquine use. In co-endemic districts, continued chloroquine use for *P. vivax* treatment likely maintains selection on *P. falciparum*, whereas in Gambella, where *P. vivax* transmission is limited and chloroquine use is minimal, *Pf*CRT K76T occurs at lower frequency (10% in Gambella versus 61.1% in all other districts, p <0.0001; **Supplementary Figure S6**). This differentiation was reflected in Jost’s D values at *Pfcrt* SNPs, where every missense SNP exceeding 0.7 came from a comparison between Gambella and another site (**Supplementary Data 2)**. In addition, PCA showed clear clustering by *Pf*CRT genotype in several districts, including Gondar, Mizan and Fentale, indicating that *Pf*CRT contributes substantially to local parasite population structure (**Supplementary Figure S7**). In contrast, K76T was fixed in Arba Minch despite presence of two distinct parasite clusters observed along PC1, indicating that population structure at this site is driven by loci outside *Pfcrt*. While the fixation of K76T is consistent with long-standing chloroquine selection in this *P. vivax* co-endemic setting, the absence of emerging variants such as *Pf*K13 R622I suggests that the evolutionary forces shaping parasite populations in Arba Minch may differ from those operating in northern Ethiopia. Beyond K76T, *Pfcrt* diversity was extensive, with 10 extended haplotypes observed in more than one Ethiopian isolate (**Supplementary Figure S8**).

**Figure 3.**
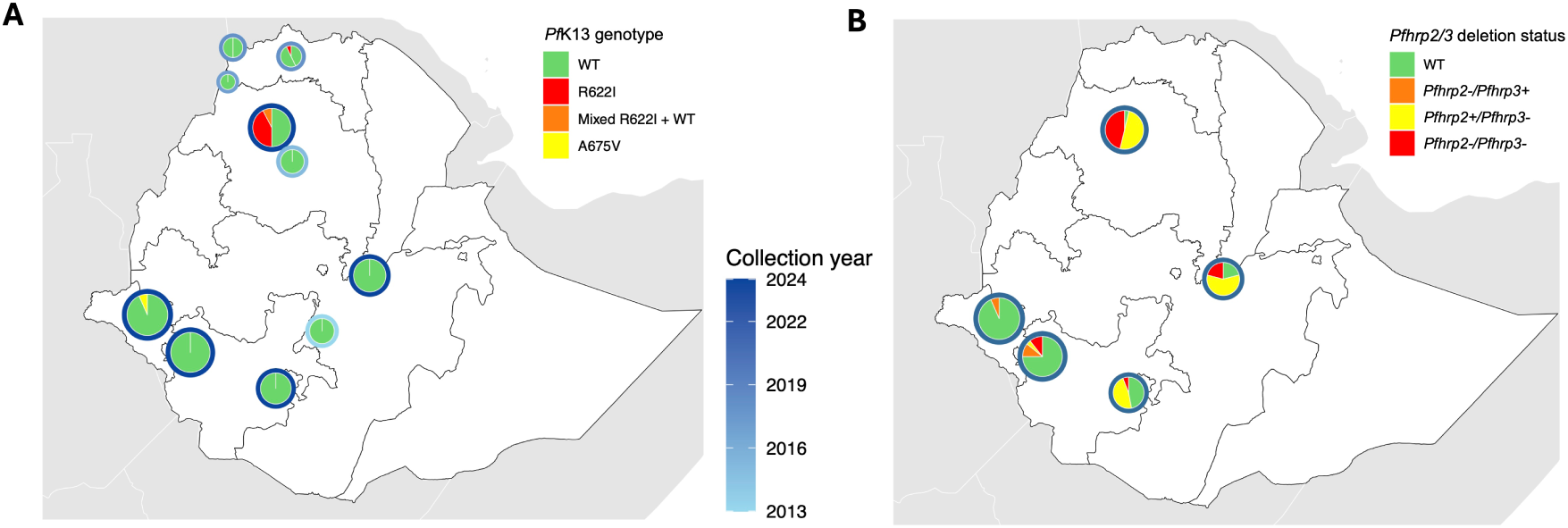
Distribution of *Pf*K13 variants and *Pfhrp* deletions. (**A**) Distribution of *Pf*K13 variants and (**B**) *Pfhrp2/3* deletions in Ethiopian isolates analysed in this study. Outer ring of pie charts indicates the sample collection year.

We next evaluated variation at loci implicated in susceptibility to dihydroartemisinin and ACT partner drugs. The recently described *Pf*PX1 PIN haplotype was present in 5.7% (6/105) of monoclonal Ethiopian isolates, all from the southwestern sites of Gambella (n=4) and Mizan (n=2), and all collected in 2024. The haplotype was absent from older Ethiopian genomes (**Supplementary Figure S9**). All isolates harbouring the PIN haplotype also carried the D384A-PIN haplotype [35]. Interestingly, both isolates carrying the *Pf*K13 A675V variant also harboured the *Pf*PX1 D384A-PIN haplotype. Allele frequencies of missense variants in known and putative drug resistance genes are provided in **Supplementary Data 3**.

Beyond adaptation affecting antimalarial treatment efficacy, we also observed widespread variation at loci associated with diagnostic evasion (**Figure 3B**). *Pfhrp2*/3 deletions were common across Ethiopia, with distinct structural patterns. *Pfhrp2* deletions encompassed *Pfhrp2* and the subtelomeric end of chromosome 8, whereas *Pfhrp3* deletions occurred through recombination between chromosomes 13 and 11 (**Supplementary Figure S10**) [36]. Double *Pfhrp2*/3 deletions were most frequent in Gondar (46.1%, 12/26), followed by Fentale (21%, 4/19), Mizan (10.7%, 3/28) and Arba Minch (5.8%, 1/17), and were absent from Gambella. Single *Pfhrp3* deletions were detected across four districts, with the highest prevalence in Gondar (50%, 13/26), whereas *Pfhrp2*-only deletions were restricted to Mizan (10.7%, 3/28) and Gambella (6.6%, 2/30) [11,37]. Notably, dual *pfhrp2/3* deletions co-occurred with the R622I variant within the same parasite genome in 46.2% (6/13) of R622I-positive isolates or 23.1% (6/26) of Gondar isolates.

### Genomic signatures of selection

Extended Haplotype Homozygosity (EHH) analysis revealed strong directional selection surrounding *Pf*K13 R622I, with the derived allele exhibiting markedly greater haplotype homozygosity than the ancestral allele (**Figure 4A**). However, the locus was not significant in genome-wide iHS scans (**Figure 4B-C**), likely reflecting the reduced power of iHS to detect selection at low frequency variants [38]. Consistent with this, the 10 kb window containing *Pfk13* was identified as under selection in Ugandan genomes (**Figure 4D**, **Supplementary Data 4**), where A675V (18.87%) and C469Y (20.25%) occur at higher frequencies than R622I (8.23%) in Ethiopia. Genome-wide scans identified two major classes of loci under selection (**Supplementary Data 4-11**). First, strong signatures were detected at established drug resistance loci, including *Pfdhps*, *Pfaat1*, *Pfmdr1*, *Pfcrt* and *Pfpx1* (**Figure 4D**). Signals at *Pfdhps* were widespread across countries and regions, consistent with sustained sulfadoxine pressure, whereas *Pfaat1* showed evidence of selection in Ethiopian, Nigerian and Ghanaian populations supported my multiple independent methods.

**Figure 4.**
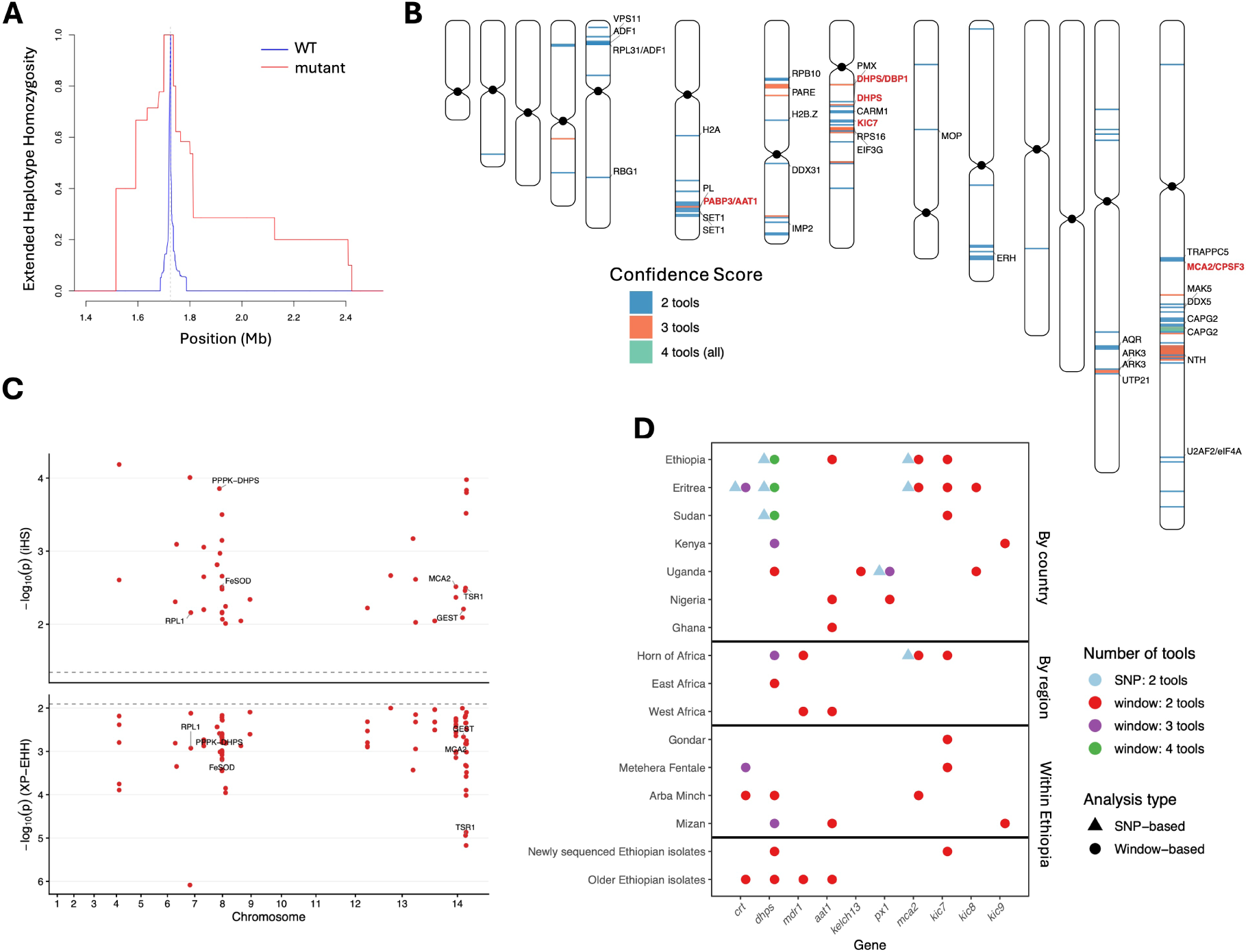
Analysis of genome-wide signatures of selective pressures in geographical and temporal comparisons of parasite populations. (**A**) Extended Haplotype Homozygosity (EHH) of the wild-type and mutant alleles in Ethiopian isolates around position 1725133 on chromosome 13, where the SNP causing the R622I variant in PfK13 is located. (**B**) Windows detected in the window-based selection analysis (iHS, XP-EHH, Tajima’s D and hmmIBD) in Ethiopian isolates, annotated per chromosome, coloured by the number of tools that detected the selection signal. Only signals detected by two or more tools are shown. Genes of interest are annotated in red. (**C**) SNP-based analysis of signals of selective pressure in Ethiopian isolates, including iHS and XP-EHH results. Dashed lines indicate the thresholds used in this composite analysis and only SNPs detected by both tools are shown here. Only genes with an assigned gene name are labelled; genes with only a gene ID are unlabelled. Annotations for all SNPs can be found in **Supplementary Data 5**. (**D**) Genes of interest showing evidence of positive selection across geographically and temporally defined population comparisons. Each point represents a selection signal detected in a gene and population. Triangles denote SNP-based analyses (iHS, XP-EHH) and circles denote window-based analyses (iHS, XP-EHH, IBD, Tajima’s D). Circle colour indicates the analysis type and the number of distinct selection tools that independently identified the signal, with blue shades for SNP-based and red/purple/green shades for window-based analyses. Only signals detected by two or more tools are shown. Populations that did not show a signal in two or more tools for any of the identified genes of interest are omitted from the Figure (Mali, Tanzania and Gambella).

Second, positive selection was detected at several genes involved in the *Pf*K13-mediated haemoglobin endocytosis pathway. The 10 kb window containing *Pfmca2* showed evidence of selection in Ethiopian parasites supported by iHS and XP-EHH. Two SNPs within *Pfmca2* (chromosome 14 positions 1,551,105 and 1,551,150) were independently identified by both SNP-based tools. Selection signals at *Pfmca2* extended beyond Ethiopia to Eritrea and Sudanese populations. Interestingly, EHH analyses suggested selection acting on the wild-type rather than the mutant allele at this locus (**Supplementary Figure S11**), with the strongest signal observed in Arba Minch. Positive selection was also detected at *Pfkic7*, a key component of the *Pf*K13-associated endocytic machinery, across Ethiopian, Eritrean, Sudanese, and the broader Horn of Africa populations, including the newly sequenced Ethiopian genomes. Additional *Pf*K13-interacting proteins showing evidence of selection included *Pfkic8* in Uganda and Eritrea and *Pfkic9* in Kenya and Mizan (**Figure 4D**).

### Genetic background of Kelch13 mutant isolates

Given the evidence for selection acting on *Pfk13* and other components of the haemoglobin-endocytosis pathway, we examined the genomic backgrounds associated with *Pf*K13 variants in the Horn of Africa. Logistic regression, incorporating the first five principal components to account for population structure, identified several loci significantly associated with R622I status, most prominently *Pf*CRT variants A220S, Q271E, N326S and R371I (**Figure 5A**; **Supplementary Data 12**), indicating that R622I is enriched within specific *Pf*CRT backgrounds. We next compared parasites carrying the ancestral R622 allele (n=146) with R622I mutants (n=13). Signals of positive selection at *Pfmca2* and *Pfkic7* were detected in the wild-type group (**Figure 5B, Supplementary Data 13-14**), with *Pfmca2* supported by hmmIBD and iHS and *Pfkic7* by Tajima’s D and hmmIBD. Together with the regional analyses, these findings indicate that adaptation within the haemoglobin-endocytosis pathway is not restricted to R622I-carrying parasites. Haplotype analysis of monoclonal isolates showed that R622I parasites from Ethiopia and Eritrea clustered together at the *Pfk13* locus (**Figure 5C**). Consistent with this, Ethiopian and Eritrean R622I isolates shared IBD across the entire gene and formed a single connected network (**Supplementary Figure 12**). At the genome-wide level, some R622I and wild-type isolate pairs showed substantial IBD (0.2-0.8), indicating intermixing of mutant and wild-type backgrounds (**Supplementary Figure S13**). Similarly, a neighbour-joining tree of the full genome placed all Ethiopian and most Eritrean R622I isolates within a single clade alongside wild-type parasites (**Figure 5D**). Restricting analyses to locations where R622I was observed (Gondar and Eritrea), R622I parasites exhibited significantly higher median genome-wide pairwise IBD fraction than wild-type parasites, except Shambuko (**Supplementary Figure S14**), consistent with local expansion of R622I bearing lineages.

**Figure 5.**
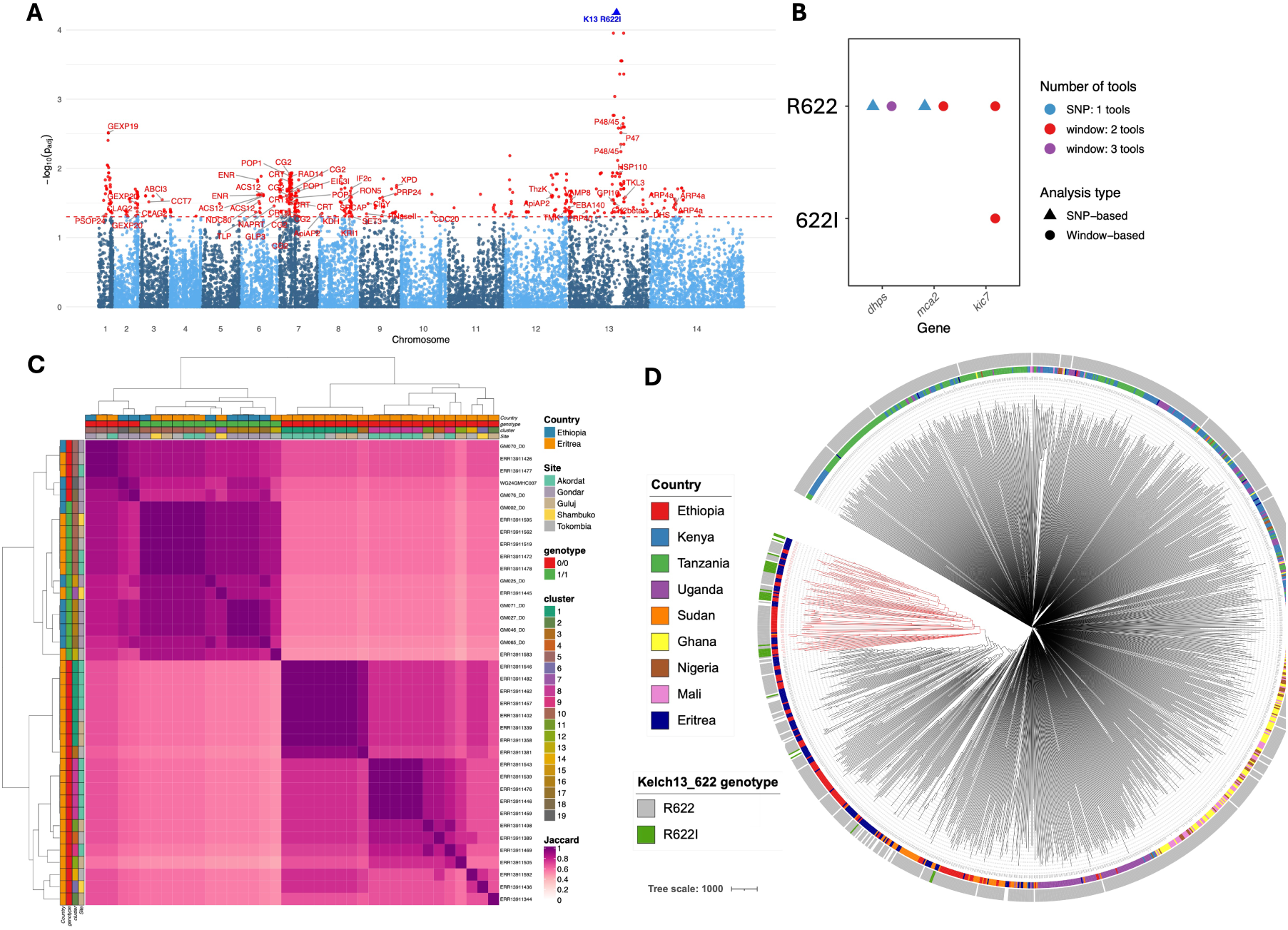
Investigation of the genomic background of *Pf*K13 mutant isolates. (**A**) Genome-wide interaction analysis between *Pf*K13 R622I and all SNPs in the Horn of Africa population (isolates from Ethiopia, Eritrea and Sudan). Each point represents a SNP, plotted by genomic position (x-axis) and significance as −log₁₀(adjusted p-value) (y-axis). Points are coloured by chromosome in alternating shades of blue, with significant SNPs (p_adj < 0.05) highlighted in red and labelled by gene name. The dashed red line indicates the significance threshold (p_adj = 0.05). The blue triangle marks the position of the *Pf*K13 R622I variant on chromosome 13. (**B**) Genes of interest showing evidence of positive selection when comparing parasites carrying the wild-type R622 genotype against those with the R622I variant in *Pf*K13. Triangles denote SNP-based analyses and circles denote window-based analyses. Point colour indicates the analysis type and the number of selection tools supporting the signal. For window-based analyses, signals detected by only a single tool are excluded. (**C**) Heatmap presenting the pairwise Jaccard index of the full *Pffi13* gene for monoclonal isolates. The annotation bar indicates sample sites, cluster assignment, *Pf*K13 622I genotype result, and country. Clusters were defined by calculating pairwise Jaccard similarity across the region and isolates with Jaccard similarity greater than 0.95 were assigned to the same cluster. (**D**) Neighbour-joining tree constructed using all samples in this study. The inner most circular annotation indicates the country origin, while the outer circular annotation represents the *Pf*K13 R622I genotype. The clade highlighted in red contains all Ethiopia isolates and some Eritrean isolates carrying the *Pf*K13 R622I variant, together with wild-type samples.

For the A675V variant, the two Gambella isolates showed greater haplotype similarity to Uganda than Southeast Asian A675V parasites, suggesting a shared or related origin (**Supplementary Figure S15**). Six Gondar isolates carried both R622I and dual *pfhrp2/3* deletions. Pairwise IBD analysis showed no evidence that that these parasites formed a distinct high-relatedness cluster, consistent with ongoing recombination with the broader parasite population rather than clonal expansion (**Supplementary Figure S16**).

### Genetic background of candidate partner drug resistance markers: *Pf*PX1 and *Pf*CRT

The recently identified *Pf*PX1 D384A-PIN haplotype was observed in 6 Ethiopia isolates, all collected in 2024, and was absent from older Ethiopian genomes (**Supplementary Figure S9, Supplementary Data 3)**. EHH analyses revealed extended derived haplotypes surrounding the PIN associated variants, particularly L1222P, D1705N and D384A (**Supplementary Figure S17**), consistent with ongoing selection despite their low frequencies. Among isolates from Gambella, Mizan, and Uganda, PIN carriers showed no greater relatedness to one another than non-carriers, indicating distribution across diverse genetic backgrounds rather than recent clonal expansion. Consistent with this, PIN carriers were dispersed throughout the neighbour-joining tree (**Supplementary Figure S18**). However, Ethiopian and Ugandan parasites carrying the PIN haplotype and the D384A variant shared IBD across the entire gene, forming a distinct connected cluster consistent with a shared origin. A single D384A-containing isolate was located outside this cluster, but that isolate was polyclonal (**Supplementary Figure S19**).

To place these findings in the broader context of antimalarial selection, we next examined genomic backgrounds associated with variation at *Pfcrt*. Although best known for its role in chloroquine resistance, *Pfcrt* variation has also been implicated in susceptibility to ACT partner drugs, making it relevant to contemporary antimalarial selection. Within *P. vivax-P. falciparum* co-endemic settings (Gondar, Mizan and Fentale), parasites carrying the ancestral K76 allele and the chloroquine resistant 76T allele revealed distinct loci under selection (**Supplementary Data 15 and 16**). Gene-level IBD analysis further resolved two major *Pfcrt* wild-type haplotype groups, with Gondar and Fentale parasites clustering separately from those in Gambella and Mizan (**Supplementary Figure S20**). These findings indicate that *Pfcrt* haplotypes are embedded within distinct genomic backgrounds and follow different evolutionary trajectories across Ethiopian transmission settings.

### Genetic background of *Pfhrp2/3* deleted isolates

To investigate the genomic architecture of diagnostic evasion variants, we characterised haplotypes surrounding *Pfhrp2* and *Pfhrp3* deletion breakpoints. *Pfhrp3* deletions were mediated by a terminal deletion of chromosome 13 encompassing the *Pfhrp3* locus, accompanied by a recombination event with chromosome 11 resulting in partial duplication of the latter. In monoclonal isolates with this deletion, the two copies of chromosome 11 exhibited sequence divergence, suggesting either recombination with a genetically distinct parasite, resulting in the acquisition of two divergent chromosome 11 copies, or a recombination event that occurred several generations ago followed by subsequent mutations [36]. In contrast, some isolates displayed no haplotype diversity between the duplicated chromosome 11 copies, suggesting recent duplication events without subsequent recombination, indicating that these events happen regularly (**Figure 6A, Supplementary Figure S21**). In addition, we found that duplicated copies of chromosome 11 cluster together with isolates without the deletion/duplication upon calculating the Jaccard index (**Supplementary Figures S22 and S23)**, meaning that *Pfhrp3* deleted isolates are intermixing with the local population rather than being imported. Taken together, this supports the occurrence of multiple independent deletion events, rather than strains descending from a common ancestral deletion.

**Figure 6.**
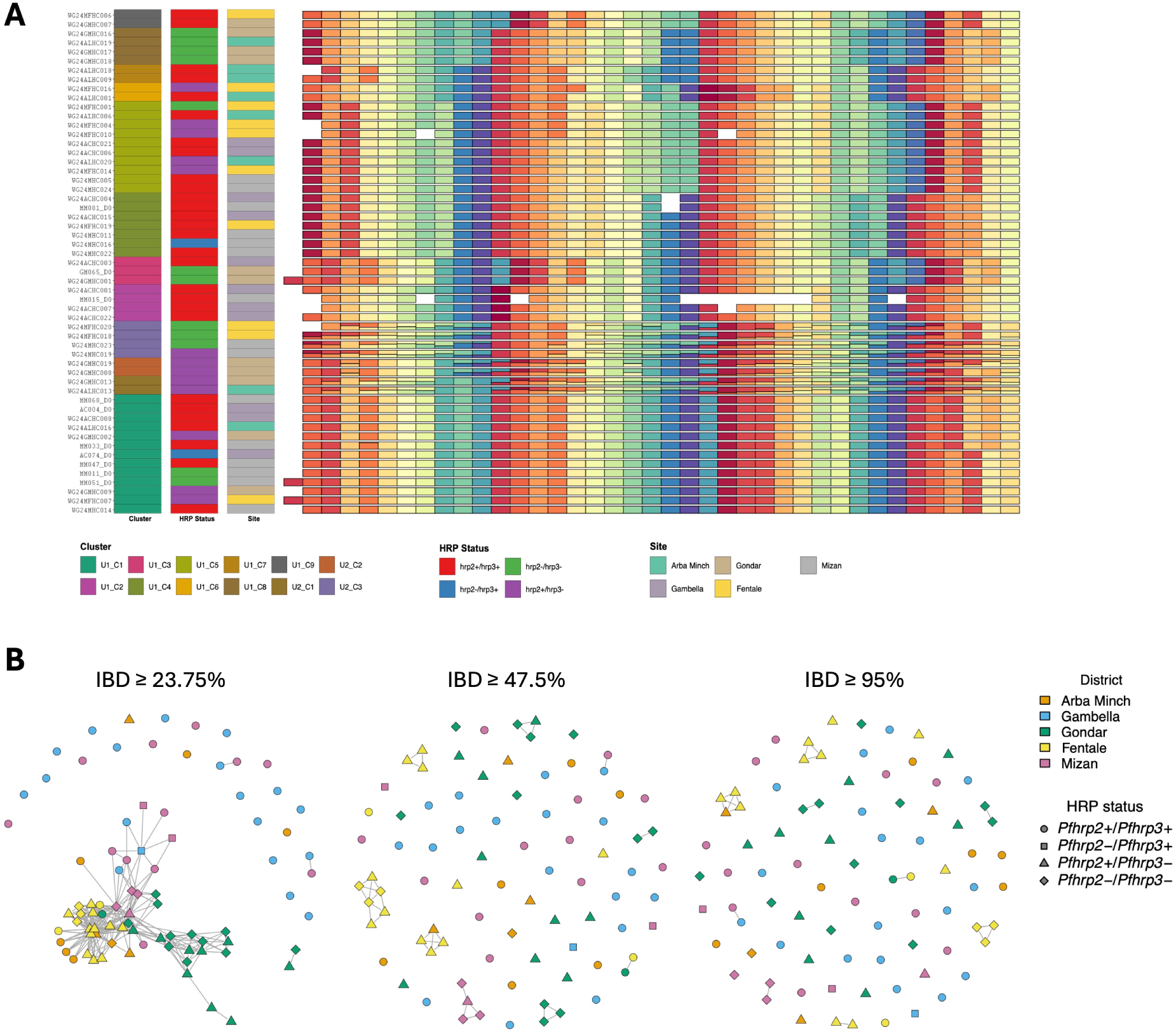
Investigation of the genomic background of *Pfhrp2/3* deleted isolates. (**A**) Haplotype diversity in duplicated region on chromosome 11 in *P5rp3* deleted isolates. Each row represents an individual isolate and each column a genomic target on chromosome 11. Within each column, cells sharing the same colour carry an identical microhaplotype at that locus; colours should be interpreted strictly per column and are not comparable across columns. The annotation bar indicates cluster assignment, *P5rp* status and sampling site respectively. Clusters were defined by calculating pairwise Jaccard similarity across the duplicate region and isolates with Jaccard similarity greater than 0.95 were assigned to the same cluster. The dataset includes both isolates with perfectly duplicated regions, characterized by a single unique haplotype, and isolates with divergent duplicate copies which exhibit two distinct haplotype structures within the region. Clusters containing only one sample were excluded from the figure. To aid visual interpretation of dominance patterns, the most frequent haplotype at each successive locus is assigned the next colour in a cycling rainbow palette. White or empty cells indicate missing data or insufficient read coverage. (**B**) Identity-by-descent (IBD) network stratified by district and *P5rp2/P5rp3* deletion status. Each node represents an Ethiopian monoclonal isolate (fws > 0.95) collected in 2024. Edges connect pairs of isolates sharing identity-by-descent (IBD) above the indicated threshold (panel titles; percentage of the genome shared IBD). Multiple panels show increasing IBD thresholds, reflecting progressively more closely related parasites, corresponding to clonal/identical (>=0.95), sibling-level (>=0.475) and related (>=0.2375) pairs. Thresholds represent successive powers of ½ (1, ½, ¼), each relaxed by 5% (i.e. multiplied by 0.95) to allow for downward bias in estimated IBD arising from missing data and genotyping error. Node colours indicate the district of sample origin. Node shapes represent pfhrp2/pfhrp3 deletion status: circles = *P5rp2+/P5rp3+* (wild type), squares = *P5rp2−/P5rp3+* (*P5rp2* deletion only), triangles = *P5rp2+/P5rp3−* (*P5rp3* deletion only), and diamonds = *P5rp2−/P5rp3−* (double deletion).

In the regions flanking *Pfhrp2* and *Pfhrp3*, EHH decayed rapidly below 0.5 in both deletion-carrying and wild-type parasites (**Supplementary Figure S24**), providing no evidence of a hard selective sweep at either locus in the 2024 Ethiopian isolates sequenced in this study. Applying the composite selection framework to comparisons stratified by *Pfhrp2/3* deletion status (*Pfhrp2*+/*Pfhrp3*+, *Pfhrp2*-/*Pfhrp3*-, and *Pfhrp2*+/*Pfhrp3*-) identified a number of windows and SNPs under positive selection within groups and in comparisons between groups (**Supplementary Data 17** and **18**). Notably, *Pfkic7* was again detected as under selection across all deletion groups, reinforcing its role as a recurrent target of selection in the Horn of Africa.

Isolates carrying one or both *Pfhrp* deletions showed higher pairwise IBD than wild-type isolates from the same district and collection year in both Mizan and Arba Minch, reaching significance in Mizan (**Supplementary Figure S25**). This is consistent with recent local expansion of deletion-bearing lineages. The *Pfhrp2-/Pfhrp3+* isolates from Mizan and Gambella isolates shared their highest IBD with *Pfhrp2+/Pfhrp3+* isolates from same districts, indicating emergence on local parasite backgrounds (**Figure 6B).**

## Discussion

This study provides a comprehensive characterization of *P. falciparum* population structure and adaptive evolution in Ethiopia using WGS of recently collected isolates. We show that Ethiopian parasite populations are highly structured both within country and relative to the wider African continent, while simultaneously exhibiting genetic connectivity with neighbouring Horn of Africa populations. Within this evolutionary framework, we detected evidence of ongoing adaptation at loci associated with drug resistance, including recent positive selection surrounding *Pf*K13 R622I, recurrent selection across multiple components of the *Pf*K13-mediated haemoglobin endocytosis pathway, the emergence of the *Pf*PX1 D384A-PIN haplotype, and widespread *Pfhrp2/3* deletions occurring on diverse local genetic backgrounds. Together, these findings suggest that parasite evolution in the Horn of Africa is increasingly characterized by repeated local emergence, recombination and expansion of adaptive variants rather than by the spread of a single dominant lineage.

Population genomic analyses identified the Horn of Africa as a genetically differentiated regional parasite population distinct from East and West Africa [39]. By integrating newly generated Ethiopian genomes with publicly available genomes from Ethiopia, Eritrea and Sudan [11,33,40], we provide improved resolution of parasite population structure across the region and confirm substantial connectivity among parasite populations within the northern Horn of Africa. Northern Ethiopian parasites showed particularly close affinity to populations from Eritrea and Sudan, indicating that parasite movement occurs across national borders and likely reflect longstanding transmission corridors. These findings indicate the importance of considering malaria evolution at a regional scale and suggest that genomic surveillance and control efforts may benefit from greater coordination across neighbouring countries.

Despite this regional connectivity, Ethiopian parasite populations exhibited marked geographic heterogeneity, supporting the existence of multiple evolutionary zones operating at finer spatial scales, as previously suggested [41]. Northern Ethiopian populations were characterised by high frequencies of *Pf*K13 R622I, extensive *Pfhrp2/3* deletions, mixed *Pfcrt* backgrounds and multiple signatures of recent adaptation. A similar mixture of *Pfcrt* backgrounds was observed in Fentale, possibly reflecting extensive labour-associated human movement linked to irrigation schemes and commercial agriculture. In contrast, Gambella clustered more closely with East African populations and harboured distinct adaptive variants, including A675V and the *Pf*PX1 D384A-PIN haplotype. Gambella also differed ecologically from most Ethiopian sites through limited *P. vivax* transmission and correspondingly reduced chloroquine pressure, resulting in predominance of wild-type *Pfcrt* backgrounds. Arba Minch, by comparison, appeared relatively isolated from emerging ART-R dynamics and remained dominated by chloroquine-resistant *Pfcrt* lineages, likely reflecting persistent selection associated with *P. vivax* co-endemicity, confirming recent observations [41]. Together, these findings suggest that parasite movement and adaptation are shaped not only by geography but also by human mobility networks, transmission ecology and local antimalarial selection pressures, generating heterogeneous evolutionary trajectories across Ethiopia.

The coexistence of regional connectivity and local population structure provides an important framework for interpreting the emergence and spread of adaptive variants. Our findings support growing evidence that ART-R in Africa is emerging through multiple independent evolutionary events rather than through introduction of a single successful resistant lineage [9,10,13–19]. R622I, the predominant validated ART-R variant in the Horn of Africa, showed strong signatures of recent positive selection, including extended haplotype homozygosity surrounding the locus. Consistent with previous reports [32], significant associations were observed between R622I and multiple *Pf*CRT variants, indicating that the emergence of R622I within parasite populations was shaped by longstanding chloroquine selection. However, R622I-carrying parasites did not form a single genetic cluster, and analyses of IBD, local haplotype sharing and phylogenetic relationships suggest that the mutation emerged on local Horn of Africa parasite backgrounds rather than through importation of an already resistant lineage. The close affinity between northern Ethiopian, Eritrean and Sudanese parasites further suggests that R622I subsequently disseminated through an interconnected Horn of Africa transmission network. In contrast, the A675V mutation detected in Gambella appears linked to East African parasite populations and transmission networks extending toward Uganda. Together, these findings support a model in which adaptive *Pf*K13 variants emerge repeatedly within local parasite populations and subsequently spread through existing regional connectivity networks shaped by human and parasite movement.

One of the most notable findings was the evidence of positive selection across multiple components of the *Pf*K13-mediated haemoglobin endocytosis pathway. Beyond selection surrounding *Pfk13* itself, we identified signals at *Pfmca2*, *Pfkic7*, *Pfkic8* and *Pfkic9* across multiple African populations. These findings are biologically plausible given the established roles of these proteins in haemoglobin endocytosis and artemisinin susceptibility. *Pf*KIC proteins and *Pf*MCA2 localize to the same cellular compartment as *Pf*K13, and disruption of *Pfmca2* and *Pfkic7* reduces artemisinin susceptibility *in vitro* [12,20]. Previous mutagenesis studies did not identify resistance-conferring mutations in several *Pf*KIC proteins [42]; however, many polymorphisms observed in our dataset were not evaluated, and their phenotypic effects may depend on genomic background [43]. Notably, signatures of selection at *Pfmca2* and *Pfkic7* were also observed among parasites lacking R622I, and in the case of *Pfmca2,* exclusively so. Together, these observations suggest that adaptation within the endocytosis pathway extends beyond validated *Pfk13* resistance lineages and supports an increasingly polygenic model of artemisinin response, whereby selection acts on multiple interacting components of the same biological pathway rather than on *Pfk13* alone.

Adaptive evolution may also be affecting ACT partner drug susceptibility. The recently described *Pf*PX1 PIN haplotype, associated with reduced susceptibility to both dihydroartemisinin and lumefantrine, was detected exclusively among Ethiopian isolates collected in 2024 and was absent from older genomes, suggesting recent emergence or introduction. Despite occurring on diverse genome wide backgrounds, PIN-haplotype parasites shared extensive local identity with Ugandan parasites, suggesting a shared evolutionary origin followed by dissemination through recombination rather than clonal expansion. The observation that both A675V-carrying parasites also harboured the *Pf*PX1 D384A-PIN haplotype further raises the possibility that partner-drug resistance backgrounds may facilitate the persistence or spread of specific *Pf*K13 variants, as has been proposed for C469Y in Uganda [35]. Although the clinical significance of these associations remains uncertain, the emergence of the *Pf*PX1 PIN haplotype warrants close surveillance given the central role of artemether-lumefantrine across Africa.

The evolutionary importance of *Pfcrt* in Ethiopian extends beyond its historical role in chloroquine resistance. Geographic variation in K76T closely mirrored patterns of *P. vivax* co-endemicity and chloroquine use, consistent with previous observations [32]. While mixed *Pfcrt* haplotypes in northern Ethiopia and Fentale contributed substantially to local population structure, Arba Minch and Gambella were largely dominated by single *Pfcrt* haplotypes and exhibited distinct evolutionary trajectories. These findings suggest that historical chloroquine selection has generated heterogeneous genomic backgrounds that continue to shape contemporary parasite evolution, including the emergence of newer adaptive variants such as R622I. Increasing evidence that *Pfcrt* variation modulates susceptibility to ACT partner drugs further highlights the continued relevance of these legacy drug selected backgrounds in the ACT era [44–47].

We also identified substantial ongoing evolution at loci associated with diagnostic evasion. *Pfhrp2/3* deletions were highly prevalent but strongly structured geographically. Dual *Pfhrp2/3* deletions reached their highest frequencies in northern Ethiopia and frequently co-occurred with R622I. In contrast, *Pfhrp2*-only deletions were restricted to southwestern Ethiopia, particularly Mizan and Gambella, differing from the historically described Ethiopian pattern in which *Pfhrp2* loss typically occurs on a *Pfhrp3*-deleted background [11,37]. Several lines of evidence support repeated local emergence rather than expansion of a single ancestral deletion lineage [11,40,48]. The chromosome 13 and 11 recombination event underlying *Pfhrp3* deletion revealed extensive haplotypic diversity, including both highly divergent and apparently recent recombination events. Moreover, deleted parasites remained embedded within local transmission networks and recombined freely with the wider population. These observations are consistent with findings from Eritrea and previous Ethiopian studies [11,27], indicating that diagnostic evasion variants arise repeatedly on diverse genetic backgrounds before undergoing local expansion. The duplicated chromosome 11 segment includes genes such as *Pf332* [49], suggesting that extra copies of this gene may confer functional consequences beyond loss of *Pfhrp3*. The restricted distribution of *Pfhrp2*-only deletions in southwestern Ethiopia further supports the possibility of additional independent emergence events.

The convergence of treatment-associated and diagnostic-evasion adaptations was particularly striking in northern Ethiopia, which emerged as the epicentre of both R622I and *Pfhrp2/3* deletions. Nearly half of R622I-positive parasites in Gondar also carried dual *Pfhrp2/3* deletions, demonstrating substantial accumulation of adaptive traits within the same parasite population. From a public health perspective, this raises concern for parasites that are simultaneously harder to detect using HRP2-based RDTs and potentially less responsive to artemisinin-based treatment. Whether this co-occurrence reflects shared selective pressures, underlying biological interactions, or simply accumulation of adaptive variants within an interconnected regional transmission network remains unclear. Nevertheless, the convergence of these traits underscores the importance of integrated genomic surveillance capable of monitoring both therapeutic and diagnostic threats simultaneously.

While these findings provide new insights into the evolutionary dynamics of *P. falciparum* in Ethiopia and the wider Horn of Africa, several considerations should be acknowledged when interpreting the results. The functional consequences of many of the variants identified within the *Pf*K13 endocytosis pathway remain unresolved and will require experimental validation. Similarly, although a subset of isolates was linked to therapeutic efficacy studies, sample sizes were insufficient for robust genome-wide association analyses of treatment outcomes. The concentration of R622I in one site in northern Ethiopia also limited our ability to fully reconstruct the diversity of genomic backgrounds on which this mutation has emerged. Finally, selective whole-genome amplification reduced sensitivity for detecting copy number variation, precluding comprehensive analysis of loci such as *Pfmdr1* and *Pfpm2/3*.

Overall, our findings indicate that malaria parasite evolution in the Horn of Africa is driven not by the spread of a single successful lineage but by repeated local emergence, recombination and expansion of multiple adaptive variants. Selection across the *Pf*K13 endocytosis pathway, emergence of candidate partner-drug resistance markers, persistence of heterogeneous *Pfcrt* backgrounds, and recurrent origins of *Pfhrp2/3* deletions collectively reveal a highly dynamic adaptive landscape. These findings highlight the need for integrated, geographically resolved genomic surveillance linked to clinical and epidemiological data to monitor the increasingly complex threats facing malaria control in Ethiopia and the wider Horn of Africa.

## Methods

### Study sites

We sampled five malaria-endemic districts in 2024 selected to capture Ethiopia’s ecological and epidemiological diversity. Maksegnit (Gondar Zuria, Amhara, near the Sudan border; hereafter “Gondar”) is a highland setting (1,750–2,600 m above sea level) with seasonal malaria transmission peaking from September-November following the main July-August rains. Metehara (Fentale, Oromia, central Ethiopia; hereafter “Fentale”) is a Rift Valley irrigation setting situated at approximately 1,000 m above sea level and centred on one of Ethiopia’s largest sugarcane plantation schemes, characterized by irrigation associated malaria transmission with seasonal peaks during September to November and substantial seasonal and permanent labour migration. Abobo (Gambella, near the South Sudan border; hereafter “Gambella”) is a lowland setting (400–600 m above sea level) with prolonged rain-linked transmission and a high malaria burden, hosting refugee populations from South Sudan as well as nomadic populations and seasonal migrant workers employed on commercial farms. Mizan-Aman (Bench-Sheko, Southwest; hereafter “Mizan”) is a mid-altitude area (1,400–1,700 m above sea level) with substantial seasonal population movement associated with agricultural activities and gold mining and has experienced a recent resurgence following a period of near-elimination. Arba Minch (Gamo Gofa, southern Ethiopia; hereafter “Arba Minch”) is a mid-altitude lacustrine Rift Valley setting with malaria endemic areas occurring primarily at 1,200–1,500 m above sea level around Lakes Abaya and Chamo. Collectively, these sites span gradients in altitude, transmission ecology, species composition, population mobility, and antimalarial selection environments. As the relative contribution of *P. vivax* generally increases with altitude in Ethiopia, co-endemic settings such as Gondar, Arba Minch, Fentale, and Mizan experience increasing use of chloroquine for *P. vivax* treatment alongside artemether lumefantrine for *P. falciparum*, whereas *P. vivax* is uncommon in the lowland Abobo. This gradient creates geographic variation in antimalarial selection pressures relevant to parasite genomic evolution.

### Sample collection

Isolates originated from two studies: a therapeutic efficacy study (TES) and molecular surveillance study (**Supplementary Figure S1**). Permissions for both studies were granted by the AHRI/ALERT Research Ethics Review Committee (AAERC) (reference numbers: PO-41-23 and PO-020-24) and the National Research Ethics Review Board (NRERB) (reference numbers: 17/152/918/23 and 17/152/542/24). For the molecular surveillance study, venous blood samples (2 mL) were collected in EDTA tubes from febrile patients with microscopy-confirmed *P. falciparum* mono-species infections and a minimum of 414 parasites/mL (n = 101) between June and November 2024. Leukocyte depletion of whole blood samples was performed using 2100ff cellulose powder (Macherey-Nagel, Germany) following established protocols (14). Red blood cells were washed with physiological saline, and the resulting pellets were stored frozen until DNA extraction. For the TES study, febrile patients older than 6 months with ≥ 1,000 asexual parasites/µL, haemoglobin ≥ 7 g/dL, and who provided informed written consent were enrolled. Prior to treatment, finger-prick blood samples were collected in EDTA-coated microtainer tubes.

### Genomic DNA extraction, molecular parasite quantification and amplification

Genomic DNA was extracted from blood using the DNeasy Blood and Tissue Kit (QIAGEN, Germany). A multiplex quantitative PCR targeting the 18S rRNA gene was used to detect and quantify *P. falciparum* and *P. vivax* parasites. Mixed species infections (*P. vivax* cycle threshold value <35, corresponding to 217 parasites/mL) were excluded. Selective whole genome amplification (sWGA) followed an established protocol [50]. DNA yield after sWGA was assessed using Qubit dsDNA High Sensitivity reagents (Invitrogen, USA) to confirm successful amplification before sequencing.

### Whole Genome Sequencing, data set selection and reads processing

Whole genome sequencing was performed for 123 Ethiopian isolates on either an Illumina NextSeq 2000 at the Armauer Hansen Research Institute (AHRI), Addis Ababa, Ethiopia, or an Illumina Novaseq 6000 platform at Eurofins Genomics, Germany, generating a minimum of 3.75 million paired-end reads (150 bp reads) per sample. The term ‘isolates’ is used to indicate parasite samples from clinical infections. A total of 1477 *P. falciparum* isolates were included in the final analysis, including the Ethiopian samples sequenced in this study, along with publicly available whole genome sequences from the Malaria Genetic Epidemiology Network (MalariaGEN) and other studies [11,14,26,51–53]. Isolates were selected from three West African countries (Ghana, Mali and Nigeria), three East African countries (Uganda, Tanzania and Kenya), all available isolates from the Horn of Africa (Ethiopia, Eritrea and Sudan), and 100 isolates from India. Where large numbers of isolates were available for a given country, the most recent collections were prioritised and representation across multiple sites was ensured. Raw sequence data were quality-filtered using Trimmomatic (v0.39) with the parameters: LEADING:3 TRAILING:3 SLIDINGWINDOW:4:20 MINLEN:36. Filtered reads were then mapped to the *P. falciparum* 3D7 reference genome (version 3) using bwa mem software (v0.7.19). SNPs and short insertions and deletions were called using samtools (v1.22) and GATK (v4.1.4.1) software. Code used for data processing can be found on https://github.com/AHRI-malaria/WGS_Pf_Ethiopia_2024. Mixed genotypes were assigned based on allele frequency, with calls made when a nucleotide comprised ≥80% of the total coverage. Isolates with over 20% missingness were excluded from all downstream analyses. Isolates from Thailand and Myanmar carrying the *Pf*K13 A675V variant were included exclusively for comparative analysis of flanking haplotypes.

### Population genetic analyses

To investigate the population structure, Principal Component Analysis (PCA) was performed using *Plink* with the filtered bi-allelic nucleotide matrix. Identical or similar isolates were removed prior to the PCA, retaining only one sample from a cluster of samples with IBD > 0.95. Clonality was assessed using the F_ws_ metric, computed within each population with the moimix R package [54]. For polyclonal isolates (F_ws_ < 0.95), the dominant allele at each heterozygous site was selected for inclusion in the PCA. Jost’s D was estimated from the multi-allelic VCF and was performed using custom R scripts. The R-based Tess3r package (version 1.1.0) was used to calculate admixture based on the spatial modelling of allele sharing using geographical coordinates from sampling sites in addition to genome-wide SNP data. The optimal number of ancestries was determined across different numbers of sub-populations (K = 2,…,10) [55]. Local haplotype construction for the known and putative drug resistance genes was conducted using PathWeaver, a tool designed to reconstruct local haplotype from microbial genome, and variants were extracted with SeekDeep [56].

A multi-tiered analytical framework integrating four complementary selection metrics was developed to identify genomic regions with robust evidence of positive selection (**Supplementary Figure S26**). Window-based signals derived from iHS, XP-EHH, hmmIBD and Tajima’s D were combined with SNP-level outputs from iHS and XP-EHH, with each window and SNP assigned a confidence score based on the number of independent methods supporting the signal. Tajima’s D was calculated with *vcftools*, in 10,000 bp windows. The *rehh* package was used to calculate the Extended Haplotype Homozygosity (EHH) around specific SNPs using the bi-allelic VCF, which uses population-based measures of haplotype diversity within (iHS) or between (XP-EHH) populations [57]. Prior to all selection analyses, input data was pruned by removing variants with an allele frequency below 0.01 within each population and multiclonal isolates were excluded (F_ws_ < 0.95) from the entire dataset (**Supplementary Figure S27**). For iHS and XP-EHH analyses, hypervariable regions were additionally excluded [58], along with 10 kb windows falling within the top 10% of Tajima’s D scores, which are indicative of balancing selection. IBD analysis was performed using *hmmIBD* software to assess connectivity between parasites within populations by estimating the pairwise fraction of shared ancestry between genomic segments, which were inferred to have descended from a recent common ancestor [59]. Flowcharts in **Supplementary Figures 26** and **27** were generated from a text-based prompt using Claude Sonnet 4.6, following which they were edited and verified by the authors.

Genomic regions detected by multiple tools were considered the most reliable candidates of regions under selective pressure. This framework was applied to multiple comparisons: between individual countries; between African regions (Horn of Africa, West Africa, and East Africa); between recently collected Ethiopian isolates (2024) and older isolates (2013–2017); between districts within Ethiopian isolates sampled in 2024; between *Pf*K13 genotypes (R622I versus wild-type); between *Pf*CRT genotypes (K76 versus 76T); and between *Pfhrp2/3* deletion profiles (*Pfhrp*2+/*Pfhrp*3+, *Pfhrp*2-/*Pfhrp*3-, *Pfhrp*2+/*Pfhrp*3-; this analysis excluded *Pfhrp*2-/*Pfhrp*3+ isolates since the sample size was low (n=5)). A list of genes of interest was assembled afterwards.

Flanking haplotype analysis was performed on monoclonal samples from Gondar and Eritrea samples. The *Pfk13* region was divided into 200 bp windows and local haplotypes were constructed using PathWeaver. Samples and targets with high proportion of missing data were excluded from the analysis. Haplotype similarity between the samples was compared using the Jaccard index, while hmmIBD was used to estimate the genetic relatedness. A logistic regression model adjusted for the top five principal components was used to identify SNPs associated with the *Pf*K13 R622I variant, as previously described [58].

### HRP deletions analysis

Local haplotype construction on these sequenced genomes was conducted using PathWeaver [36] (https://github.com/nickjhathaway/PathWeaver). Reads were mapped to a region of interest and then a de Brujin graph was constructed while keeping read information to inform graph traversal to prevent chimera assemblies in polyclonal isolates. The haplotypes from PathWeaver were analysed by constructing a graph for each genomic region to determine the conserved subregions that all haplotypes traversed using custom C++ scripts. The regions analysed were chromosome 5 (including pfmdr1): 929,384–988,747, chromosome 8: 1,290,239–1,387,982, chromosome 11: 1,897,151–2,003,328, chromosome 13: 2,769,916–2,844,785 [36]. The coverage results generated from these regions were calculated and normalized across isolates to account for differences in sequencing depth and variability. The resulting normalised coverage at *Pfhrp2* and *Pfhrp3* genes was then used to estimate the deletion status. In order to compare haplotype diversity in the duplicated region on chromosome 11 between isolates, the Jaccard index was calculated for each isolate comparison over the loci where both samples have sufficient data and was calculated as the number of microhaplotypes shared between sample A and sample B divided by the total number of microhaplotypes. When utilizing the Jaccard index as a distance between samples it was converted by computing 1-Jaccard index.

## Data Availability Statement

Raw sequencing data for genomes generated in this study can be found in European Nucleotide Archives (ENA, project accession PRJEB126687). Sample accession numbers and corresponding metadata can be found in **Supplementary Data 1**.

## Code Availability Statement

Custom scripts generated for the analysis presented can be found at https://github.com/AHRI-malaria/WGS_Pf_Ethiopia_2024.

## Acknowledgements

We thank the study participants from the five districts across Ethiopia for their cooperation. We also thank Melissa Conrad, Karamoko Niare and Didier Menard for providing additional metadata for publicly available genomes.

## Author contributions

L.N.V., D.F.W, M.K., F.G.T conceived and designed the experiments. D.F.W., M.K., H.A. and F.A.K coordinated sample collection and data management. D.F.W., M.A., A.S. and M.S. conducted sample processing and DNA extraction. L.N.V., M.S., D.W., Y.T., S.A., A.S. conducted selective whole genome amplification and sequencing. L.N.V., M.Y., F.M.M. and N.H. performed quality control of sequencing data, bioinformatics and statistical analyses. N.H. and N.B. provided bioinformatics analysis tools and scripts. L.N.V and M.Y. wrote the manuscript with input from N.B., S.C., B.G., N.H. and F.G.T. All authors reviewed, edited, and approved the final manuscript.

**Competing interests:** All authors declare no competing interests.

