## Supplementary Figure or Table for "Multiple adaptive trajectories shape antimalarial resistance evolution in Ethiopian *Plasmodium falciparum*"

### Supplementary Materials

**Fig S1. Map of samples by study**

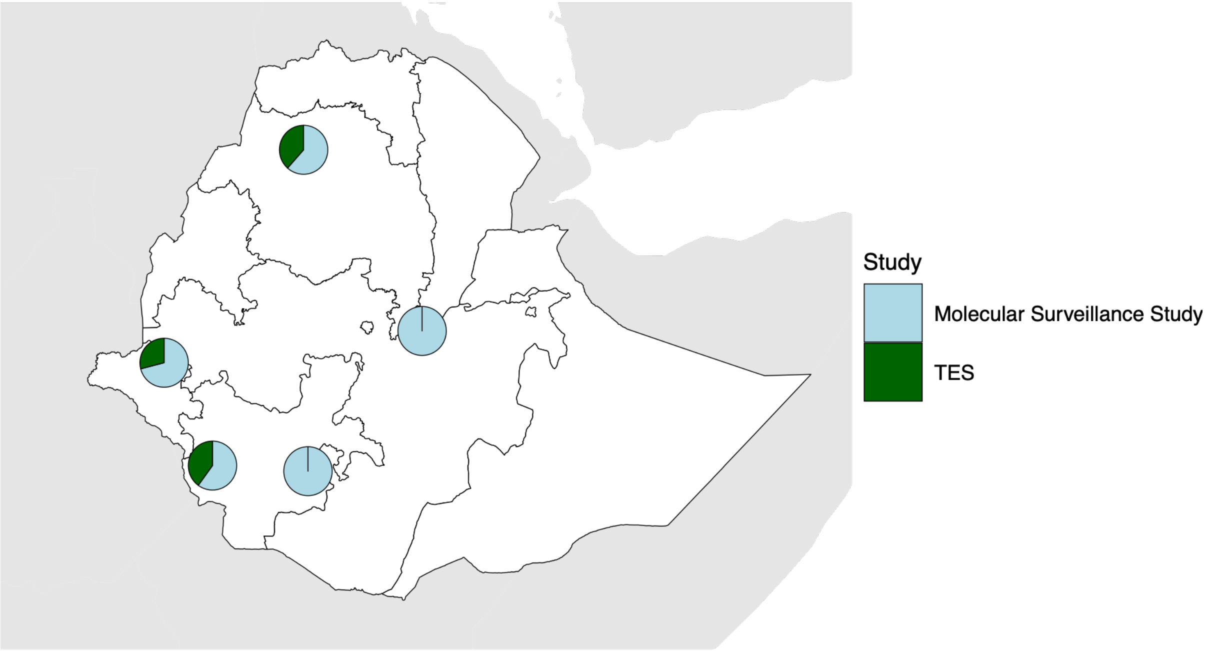

*Pie charts indicate the proportion of samples from each collection site contributed by each of the two studies included in this analysis.*

**Fig S2. Sequencing quality control metrics**

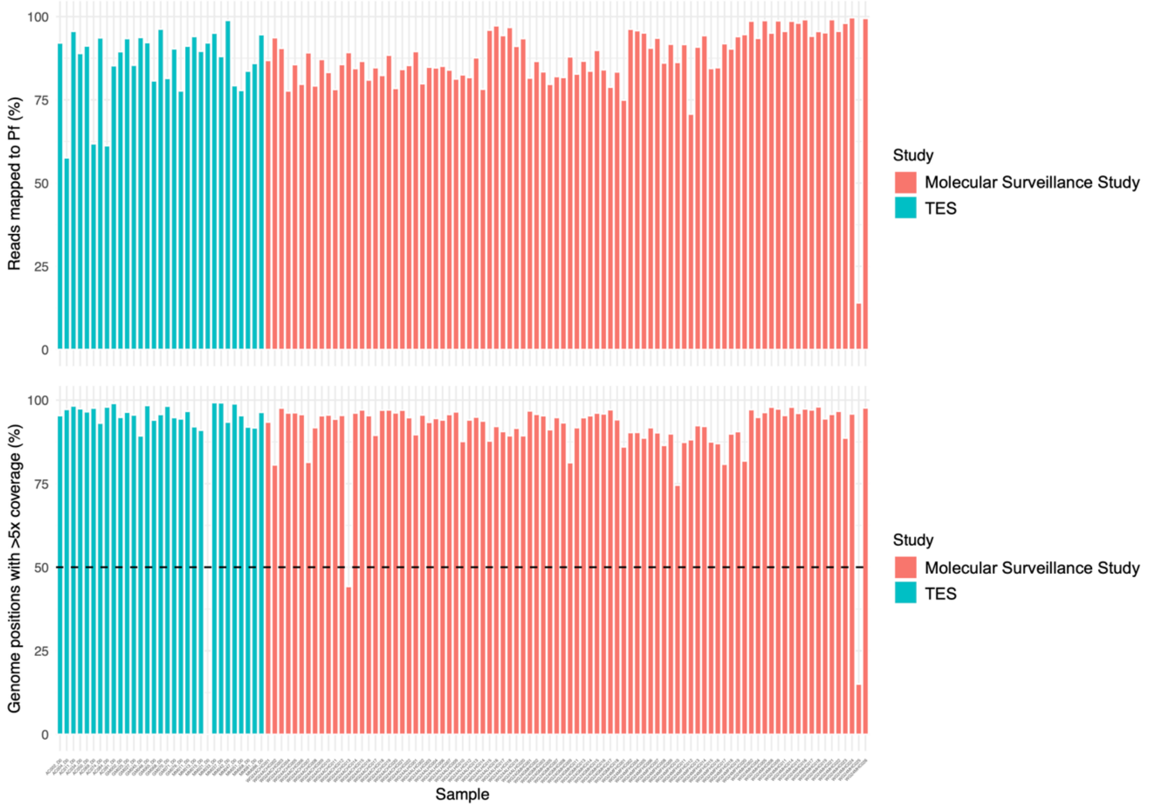

*Percentage of reads mapped (upper panel) and percentage of the genome with >5x coverage (lower panel)*

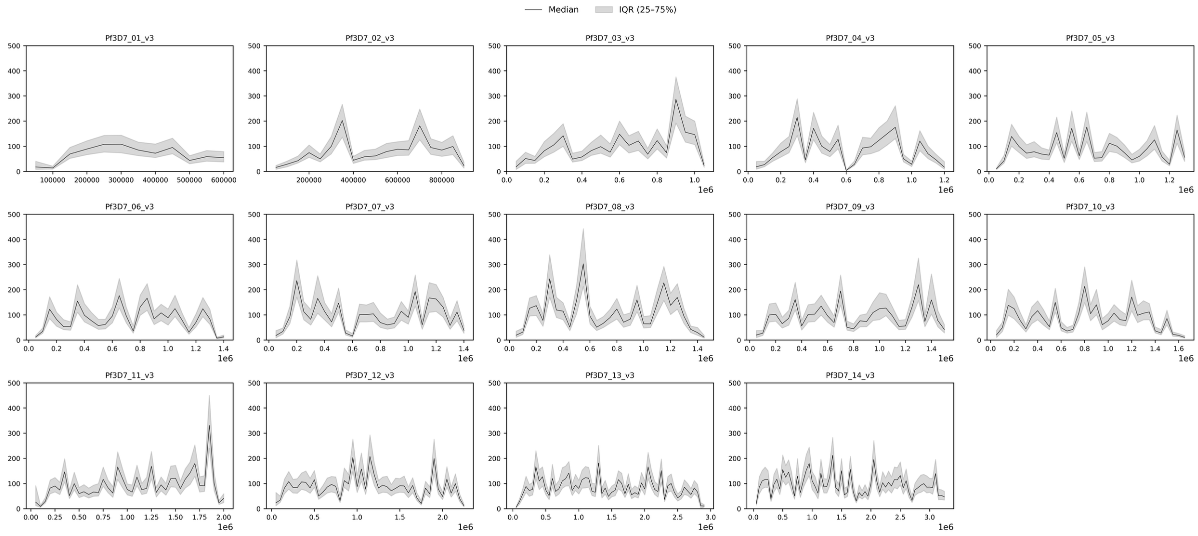

*Coverage across chromosomes in newly sequenced Ethiopian samples*

**Fig S3. Additional PCA graphs**

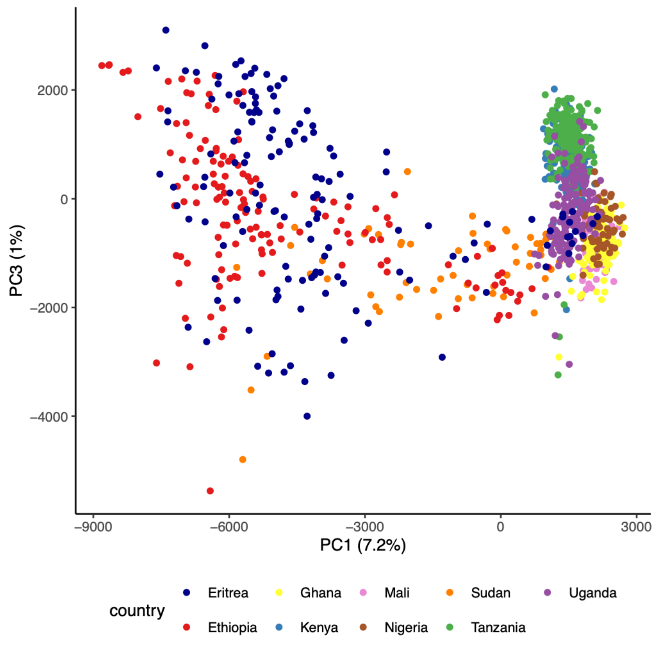

PC1 versus PC3, annotated by country

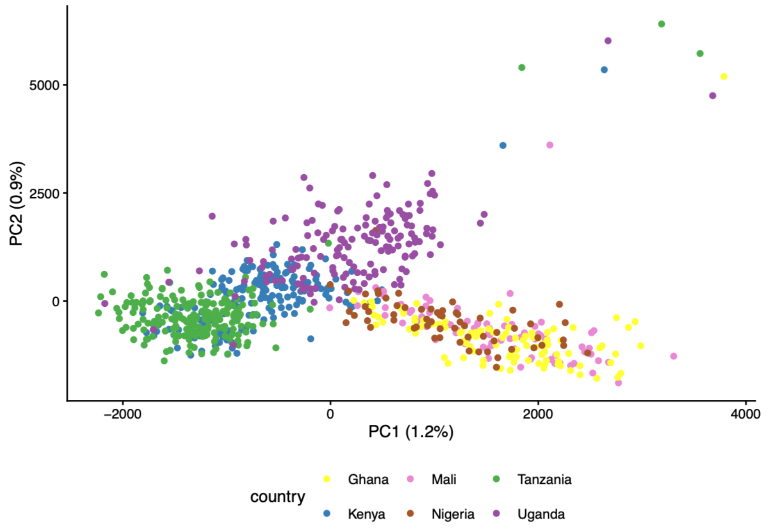

PCA of East and West African isolates, excluding Horn of Africa isolates

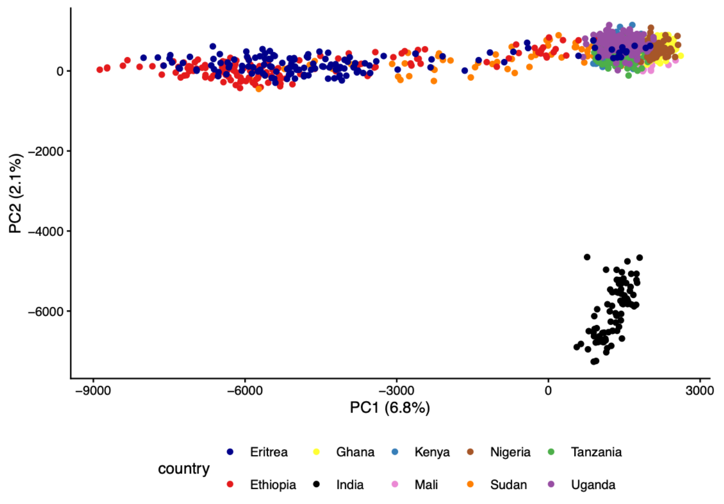

PCA including 98 Indian *Plasmodium falciparum* genomes

**Fig S4. Ancestry composition inferred by TESS3r for K = 2–10.**

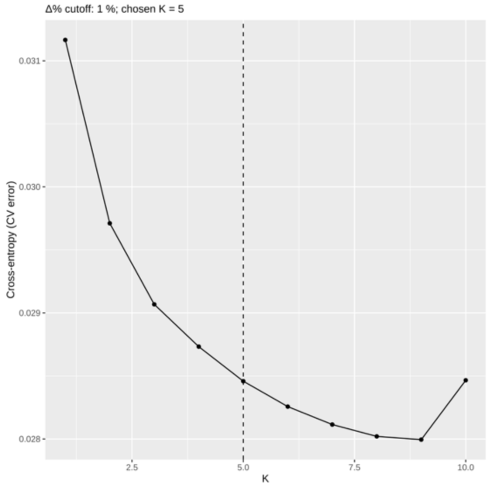

*(Top) Cross-validation error as a function of the number of ancestral populations (K). The optimal K was selected using a 1% improvement cutoff, whereby the lowest K at which the reduction in cross-validation error between successive values of K falls below 1% is chosen as optimal; this criterion identified K = 5 (dashed vertical line) as the best-supported number of ancestral populations. (Bottom) Admixture bar plots for K = 2 to K = 10, where each vertical bar represents an individual isolate and the coloured segments indicate the estimated proportion of ancestry derived from each of K ancestral populations. Samples are grouped country along the x-axis, as indicated by labels below each panel.*

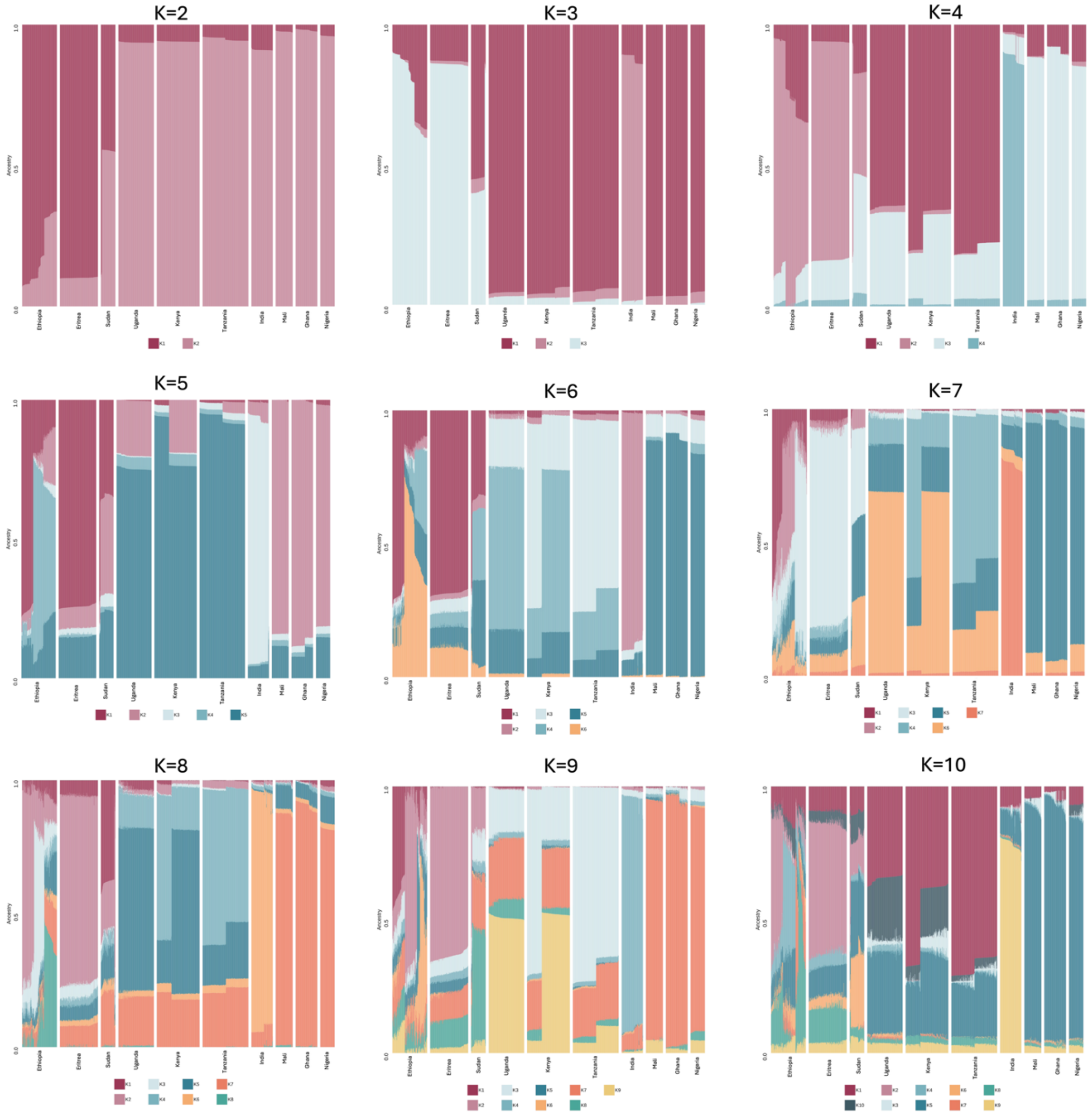

**Fig S5. PCA of all Ethiopian isolates analysed in this study and Kelch13 variant annotations**

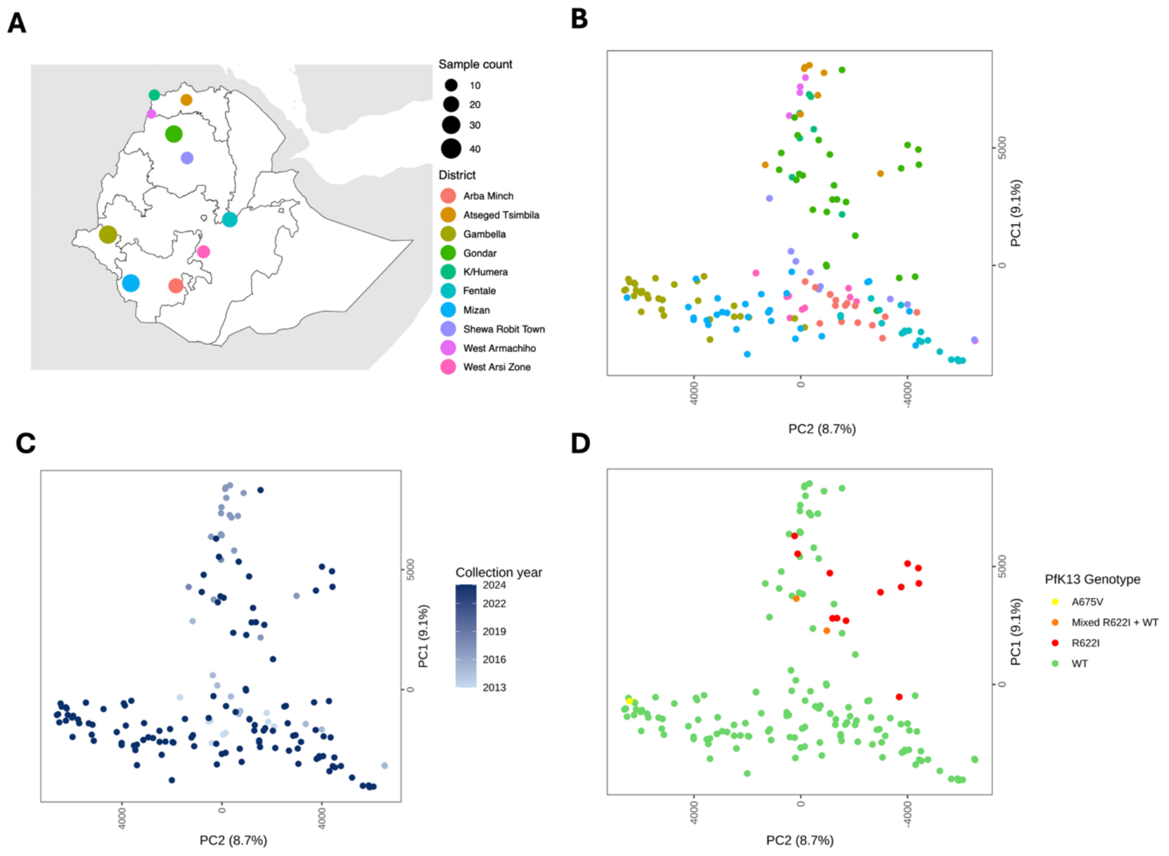

*(**A**) All Ethiopian isolates analysed in this study, including newly sequenced isolates collected in 2024, as well as older publicly available isolates. (**B**) PCA coloured by site. (**C**) PCA coloured by collection year. (**D**) PCA coloured by PfK13 genotype.*

**Fig S6. CRT variant map and annotated PCA**

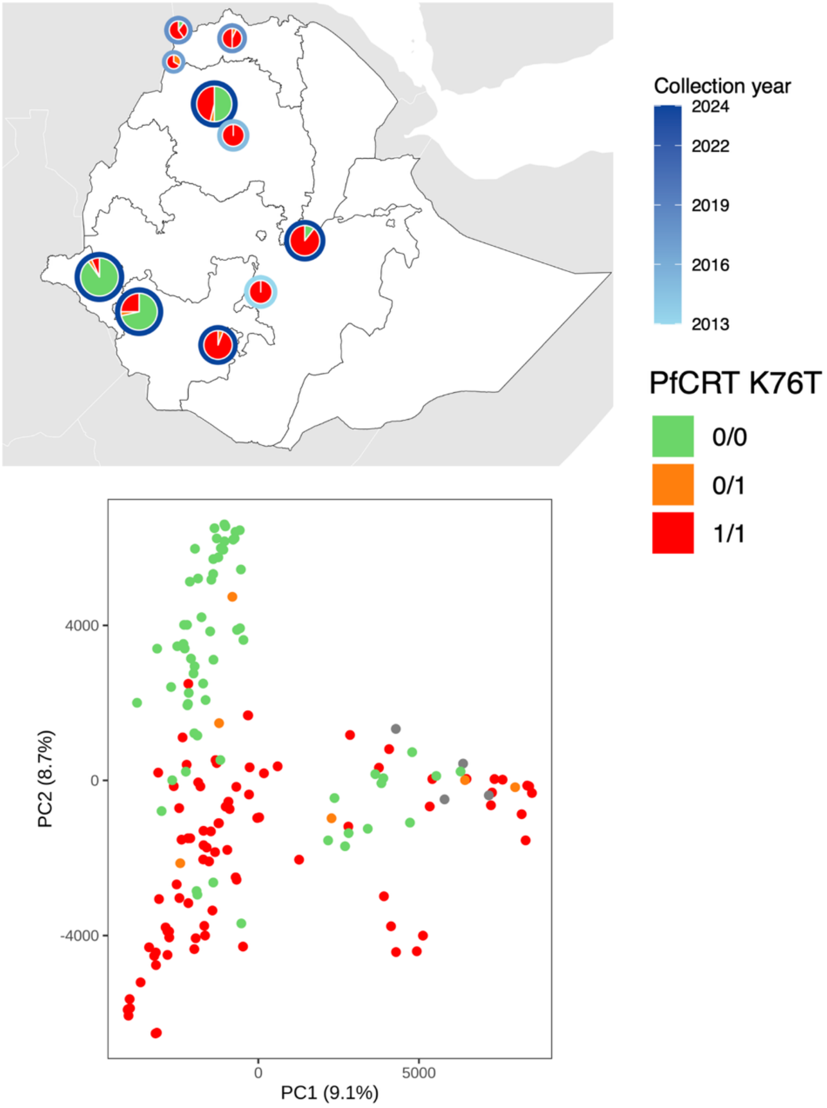

*(**A**) PfCRT genotypes in all Ethiopian samples analysed in this study. (**B**) PCA coloured by PfCRT genotype. Grey indicates the absence of a genotype call due to inadequate coverage.*

**Fig S7. Per-district PCA for Ethiopian districts, coloured by PfCRT allele at codon 76**

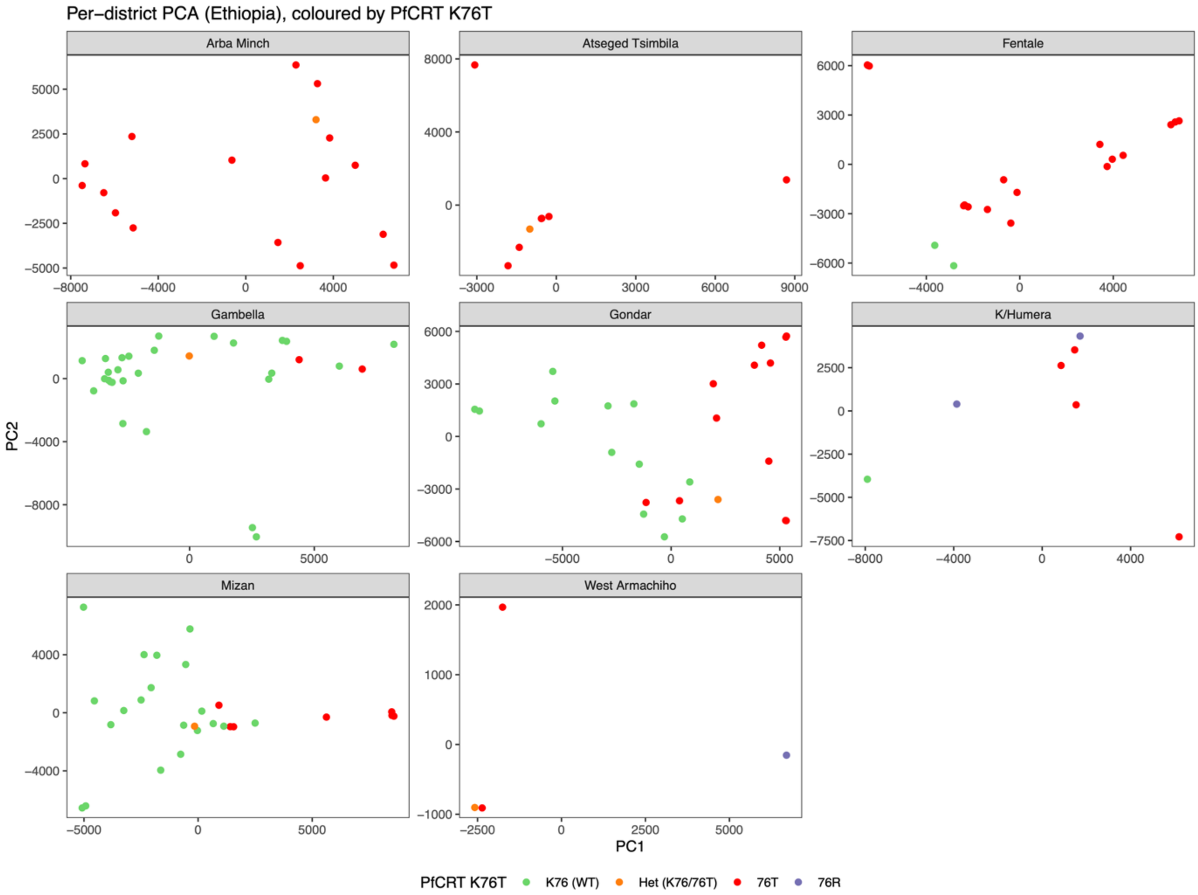

*PCA per district within Ethiopia, annotated by PfCRT allele at codon 76. Districts with only 1 allele (76T only; Shewa Robit Town and West Arsi Zone) were not included in this visualisation.*

**Fig S8. *Pf*CRT haplotypes in Ethiopia**

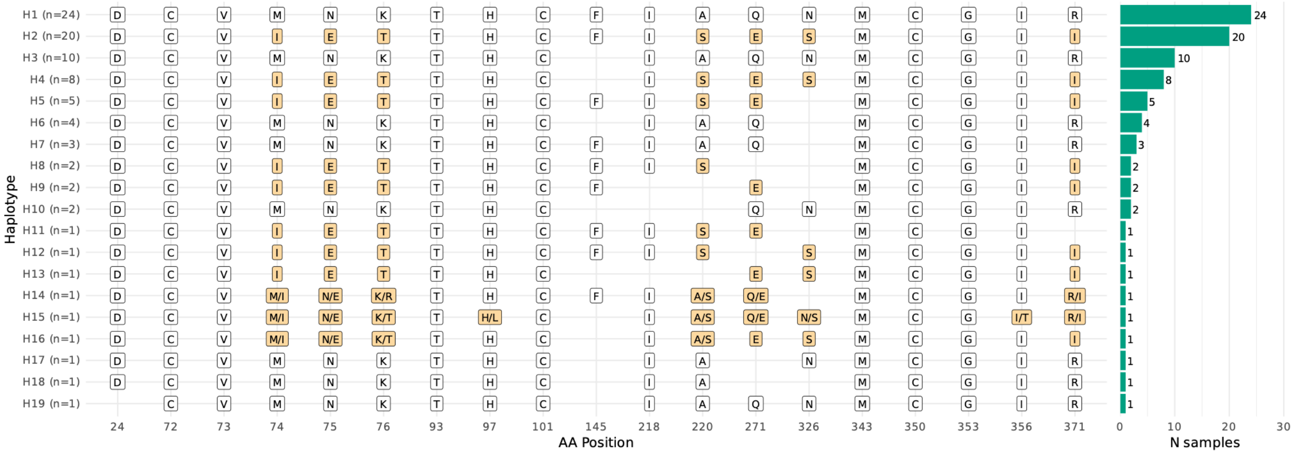

*Amino acid haplotypes reconstructed across PfCRT codons for Ethiopian isolates. Each row is a distinct haplotype, ordered by frequency, with the number of isolates carrying it given in parentheses and shown in the bar chart on the right. Residues are labelled by single-letter code; positions differing from the 3D7 reference are shaded, and unshaded cells indicate the reference allele. Slash-separated labels (for example M/I at codon 74) denote heterozygous or mixed calls within an isolate. Blank cells indicate positions with insufficient coverage to make a call. Isolates were included only if at least 80% of the genotyped positions could be called; positions failing this threshold within retained isolates are left blank.*

**Fig S9. Frequency and distribution of PX1 variants**

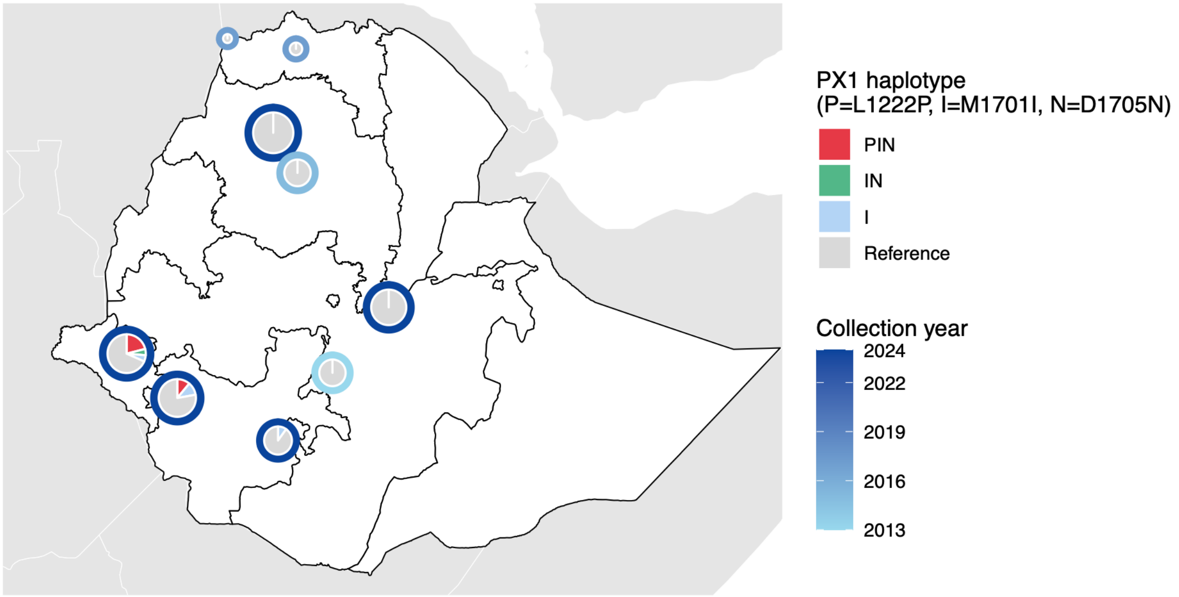

*Distribution of the PX1 PIN haplotype in monoclonal Ethiopian isolates. Outer ring of pie charts indicates the sample collection year.*

### Fig S10. Assessment of *Pf*HRP2/3 deletions

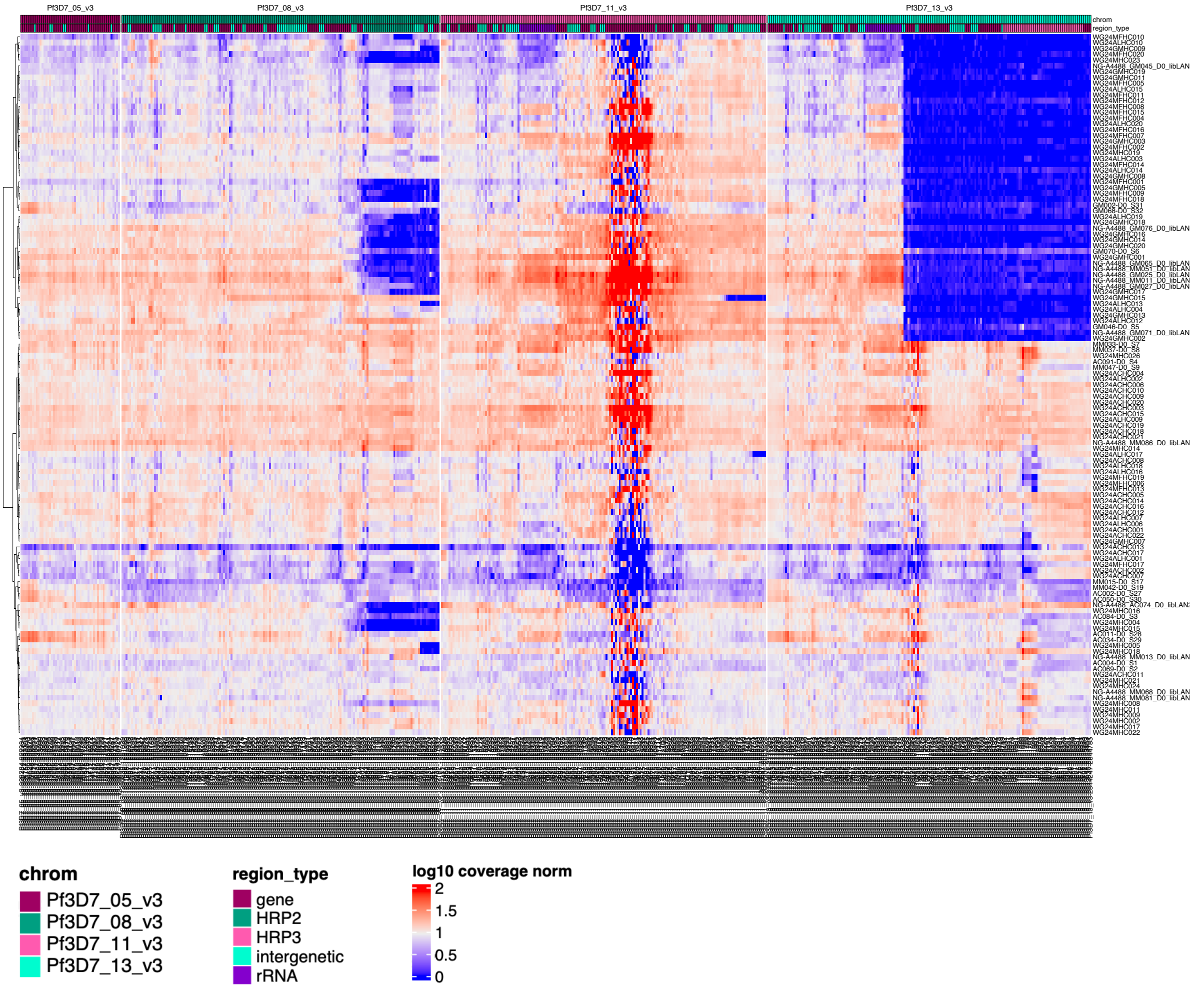

*Normalised coverage graph of regions of interest for determining PfHRP deletions on Chromosomes 5, 8, 11 and 13. Blocks of low normalised coverage (dark blue) indicate PfHRP2/3 deletions.*

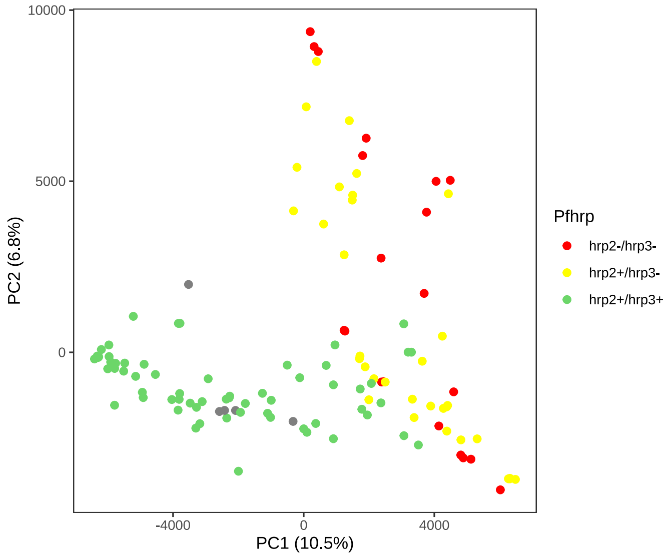

*PCA of 2024 Ethiopian isolates coloured by Pfhrp deletion status*

### Fig S11. EHH signals in *Pfmca2*

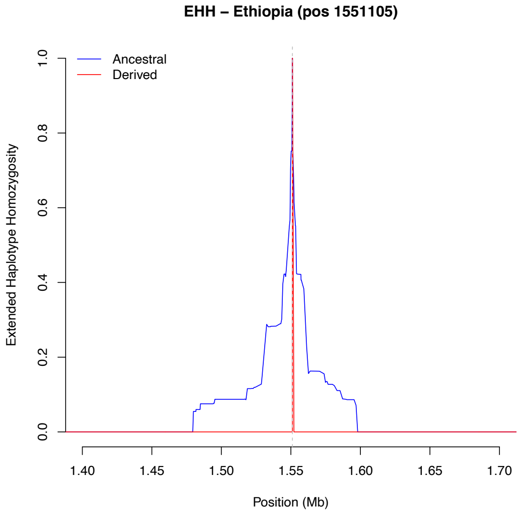

*Extended Haplotype Homozygosity (EHH) of the mutant and wild-type alleles in Ethiopian isolates around position 1551105 in Pfmca2 on chromosome 14.*

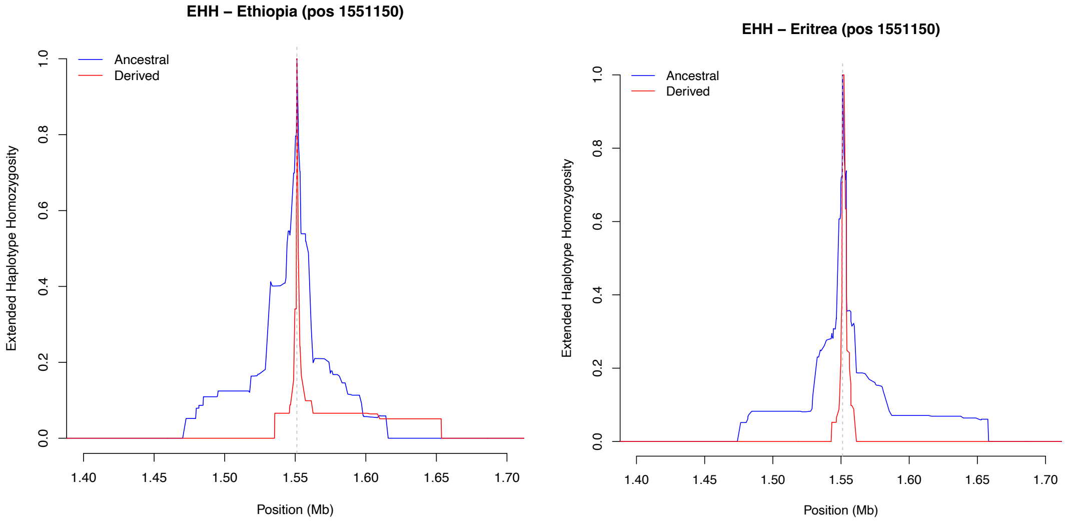

*Extended Haplotype Homozygosity (EHH) of the derived and ancestral alleles in Ethiopian and Eritrean isolates around position 1551150 in Pfmca2 on chromosome 14.*

**Fig S12. Identity-by-descent networks and haplotypes at the *Pfk13* locus.**

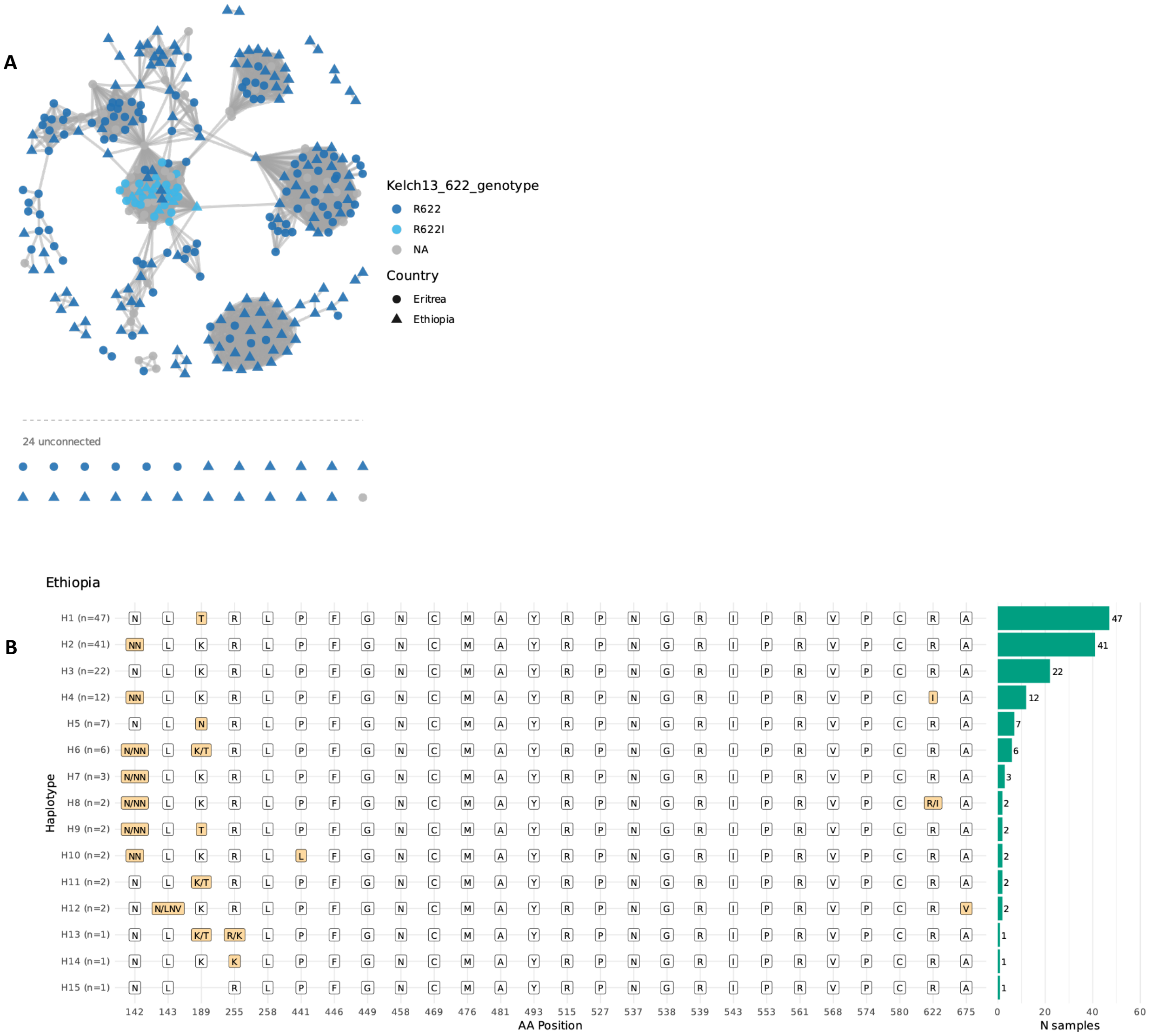

***A)*** *Each node represents one isolate (n = 323) and each edge a pair of isolates sharing a single IBD segment spanning the entire Pfk13 gene (3,249 pairs). Node colour indicates the genotype at Kelch13 codon 622 and node shape indicates country of origin (circles, Eritrea; triangles, Ethiopia). The 24 isolates sharing the full interval with no other sample are shown below the dashed line. R622I isolates from both Ethiopia and Eritrea form a single connected group, consistent with a shared origin of the mutant haplotype. For polyclonal samples, the majority allele at each site is shown.* ***B)*** *Amino acid haplotypes reconstructed across PfK13 codons for Ethiopian isolates. Each row is a distinct haplotype, ordered by frequency, with the number of isolates carrying it given in parentheses and shown in the bar chart on the right. Residues are labelled by single-letter code; positions differing from the 3D7 reference are shaded, and unshaded cells indicate the reference allele. Slash-separated labels denote heterozygous or mixed calls within an isolate. Blank cells indicate positions with insufficient coverage to make a call. Isolates were included only if at least 80% of the genotyped positions could be called; positions failing this threshold within retained isolates are left blank.*

### Fig S13. Heatmap showing Jaccard index of *Pfk13* gene and pairwise IBD fractions

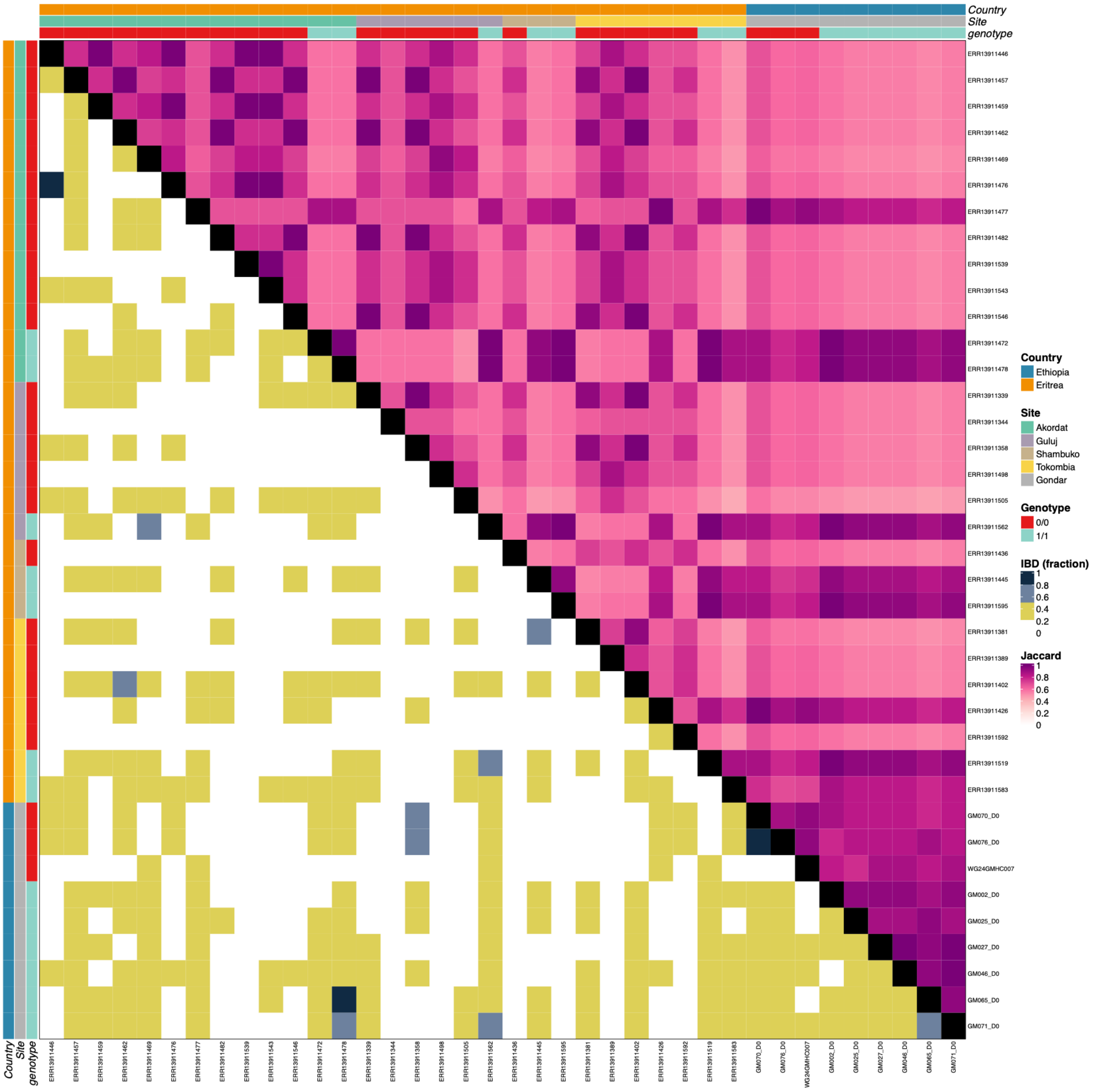

*Heatmap presenting the pairwise Jaccard index of the full Pfk13 gene for monoclonal isolates, as well as the pairwise IBD fraction of the full genome. The first colour bar indicates the country origin, the second indicates the sampling sites, and the third represents the K13 622I genotype. The upper-right half of the heatmap shows the jaccard similarity between the pairs of samples. The lower-left half displays the IBD fraction, which is grouped into four colour categories.*

**Fig S14. Pairwise IBD analysis stratified by district and R622I deletion status**

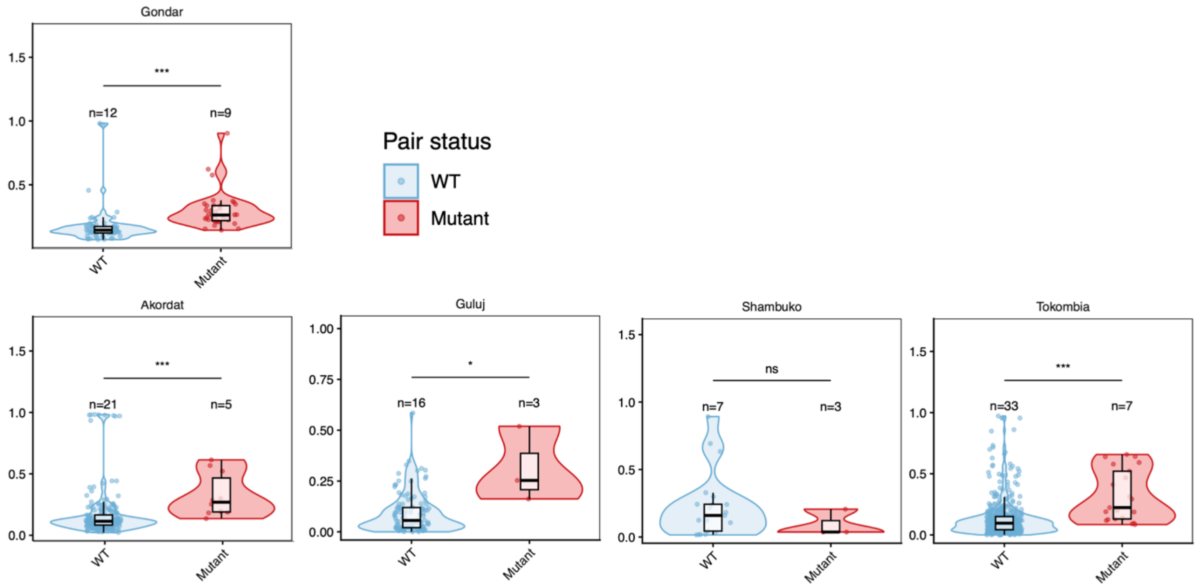

*Violin plots show the distribution of pairwise IBD (fraction of the genome shared IBD) between parasite monoclonal samples within the same district. Comparisons are restricted to pairs where both samples are from the same district and have known R622I status. Pairs are grouped as wild type (WT) or mutant (R622I). Mixed pairs (one wild type and one mutant) are excluded. Overlaid boxplots indicate the median and interquartile range, and individual points represent pairwise comparisons. Numbers above each group indicate the number of individual samples contributing to each category per district. Facets correspond to districts, with y-axis scales allowed to vary between districts. Wilcoxon rank-sum test (*p < 0.05, **p < 0.01, ***p < 0.001, ns = not significant).*

### Fig S15. Flanking haplotypes around *Pf*K13 A675V

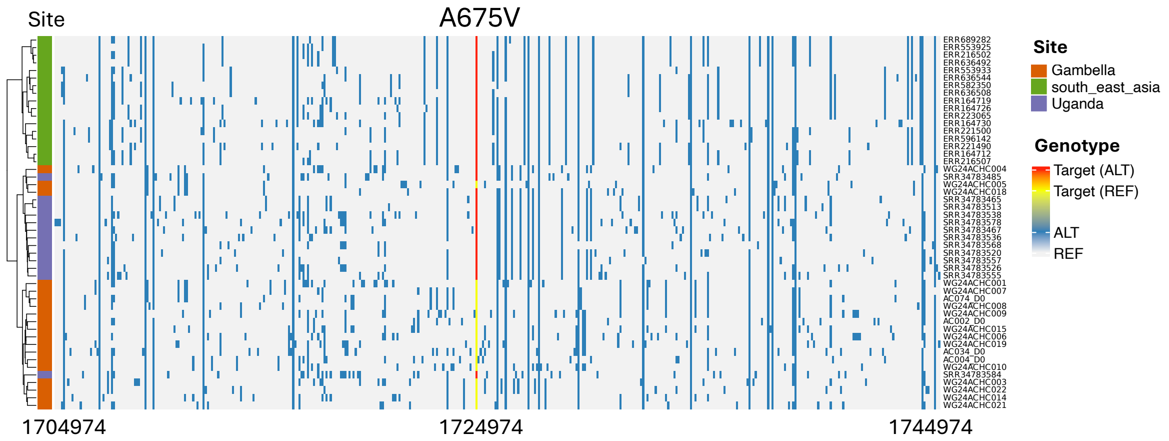

*Flanking haplotype structure around the PfK13 A675V locus (position 1,724,974) in all isolates Gambella and Ugandan and Southeast Asian isoaltes with the A675V variant. Each row represents a sample and each column a variant site, with samples ordered by hierarchical clustering. Light grey indicates the reference allele, blue indicates the alternate allele, and the A675V site is distinguished with separate colours (yellow = reference, magenta = alternate). Polyclonal samples were excluded for flanking haplotype analysis.*

**Fig S16. Pairwise IBD networks co-occurrence of R622I and HRP deletions**

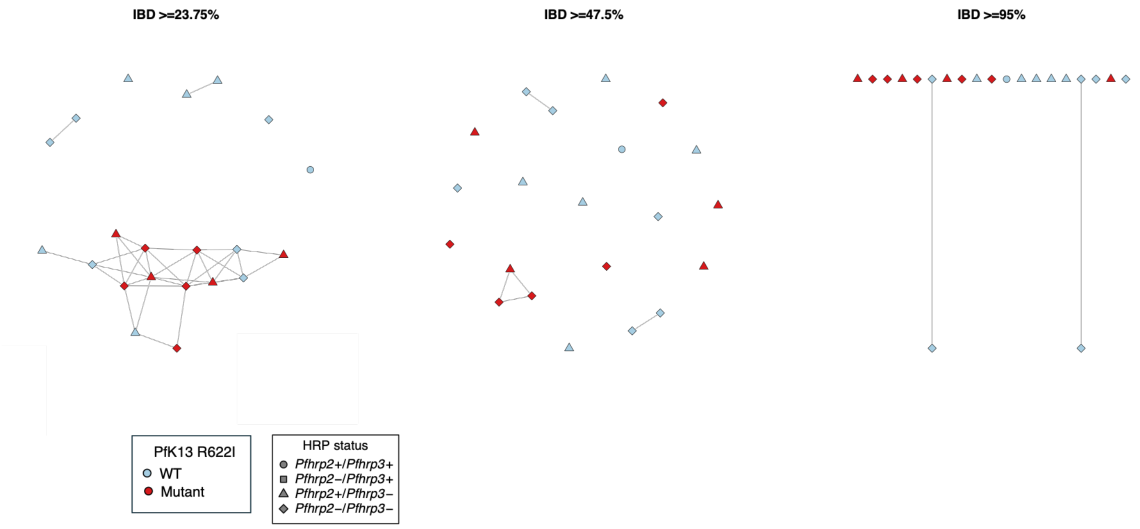

*Genome-wide identity-by-descent (IBD) network of monoclonal isolates from Gondar, stratified by PfK13 R622I deletion status and HRP2/3 deletion status.*

**Fig S17. Investigation of EHH decay around *Pf*PX1 variants**

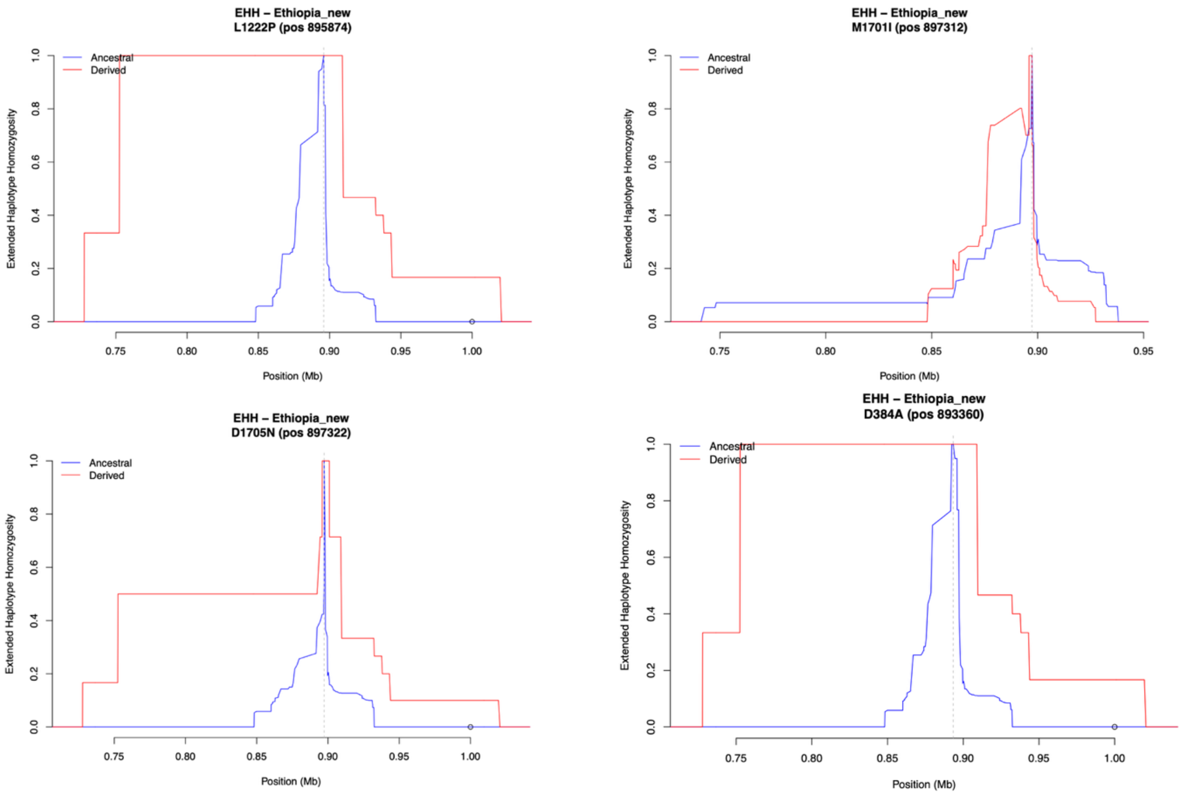

*(**A**) Extended Haplotype Homozygosity (EHH) of the derived and ancestral alleles in 2024 Ethiopian isolates around positions 895874, 897312 and 89322 on chromosome 7, where the SNPs causing the PIN haplotype in the PfPX1 gene are located, as well as a recently identified SNP at position 893360 causing the D384A variant.*

**Fig S18. Investigation of genetic background in *Pf*PX1 PIN isolates**

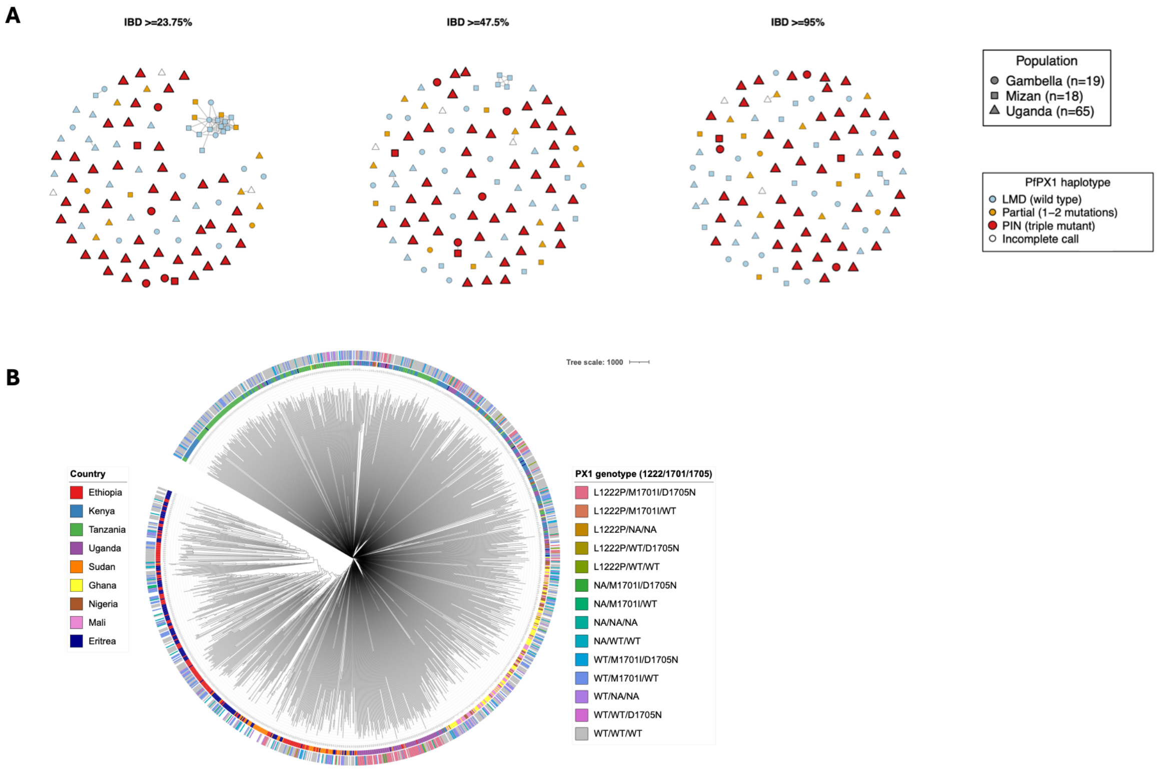

*(**A**) Genome-wide pairwise identity-by-descent networks showing isolates from Gambella and Mizan, districts in Ethiopia where the PIN haplotype was observed, as well as Ugandan isolates, the population in which the PIN haplotype was identified as under selection (Niare et al. Nat. Medicine, 2026). (**B**) Neighbour-joining tree of all African isolates in the analysed dataset. The inner ring represents the country of origin, while the outer ring represents the PX1 haplotype at amino acid positions 1222, 1701, and 1705.*

**Fig S19. Identity-by-descent networks at the *Pfpx1* locus**

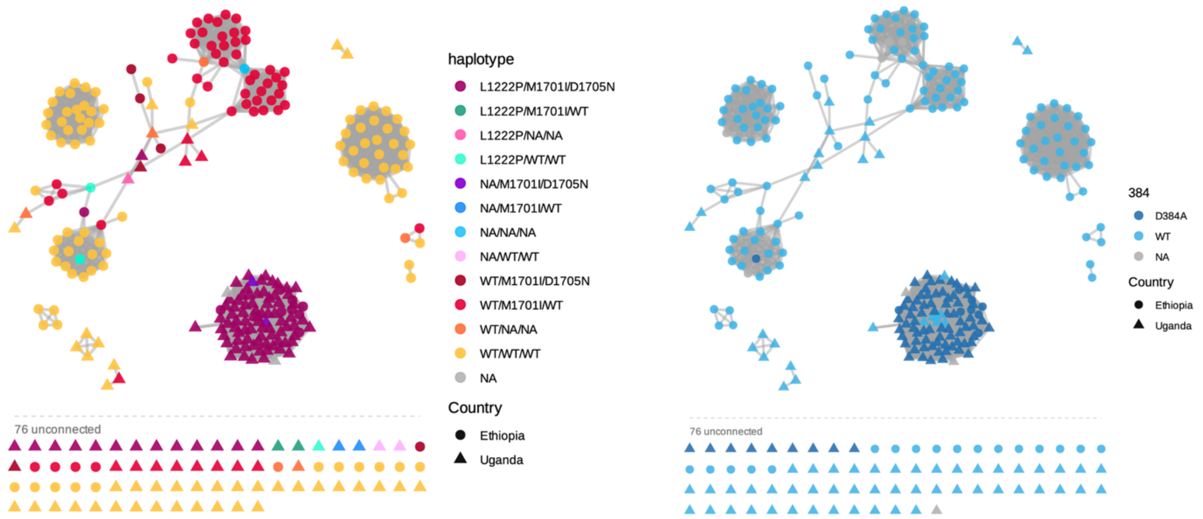

*Each node represents one isolate and each edge a pair of isolates sharing a single IBD segment spanning the entire px1 gene. Node shape indicates country of origin (circles, Ethiopia; triangles, Uganda). In the left panel, node colour indicates the multi-locus px1 haplotype, including the PIN haplotype and the wild-type and intermediate combinations listed in the key. In the right panel, node colour indicates the genotype at codon 384 (D384A, wild type, or NA where the site could not be called). The 76 isolates sharing the full interval with no other sample are shown below the dashed line in each panel. For polyclonal samples, the majority allele at each site is shown.*

**Fig S20. Identity-by-descent networks at the *Pfcrt* locus**

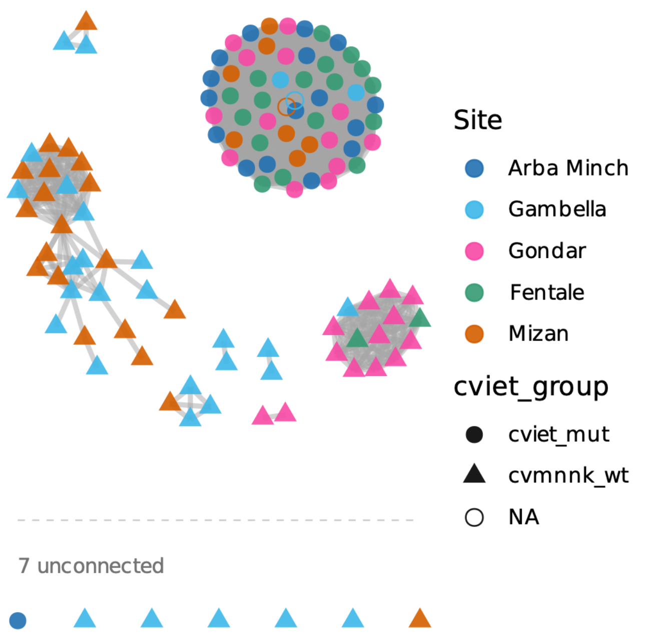

*Each node represents one isolate (n = 120) and each edge a pair of isolates sharing a single IBD segment spanning the entire Pfcrt gene (1,783 pairs). Node colour indicates collection site (Arba Minch, Gambella, Gondar, Fentale, Mizan) and node shape indicates the Pfcrt haplotype (cvmnnk wild type, cviet mutant, or NA where the haplotype could not be called). The 7 isolates sharing the full interval with no other sample are shown below the dashed line.*

### Fig S21. Haplotype diversity related to *Pf*HRP3 deletions

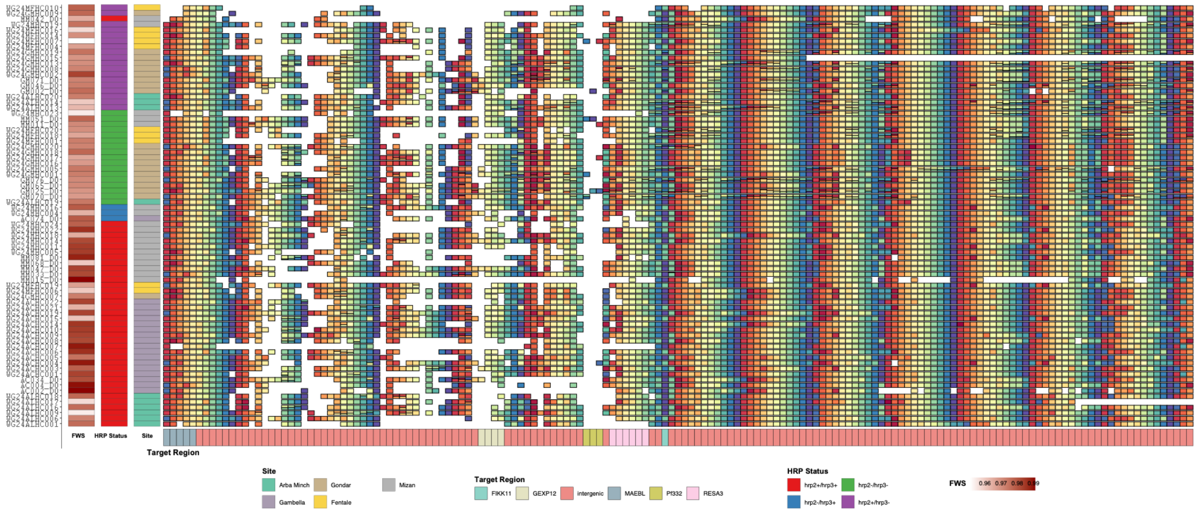

*Diversity in regions of interest for assessing PfHRP3 deletions. Each row represents an individual isolate and each column a genomic target on chromosome 11. Within each column, cells sharing the same colour carry an identical microhaplotype at that locus; colours should be interpreted strictly per column and are not comparable across columns. To aid visual interpretation of dominance patterns, the most frequent haplotype at each successive locus is assigned the next colour in a cycling rainbow palette. White or empty cells indicate missing data or insufficient read coverage.*

**Fig S22. Jaccard index between isolates with 1 unique copy of chromosome 11**

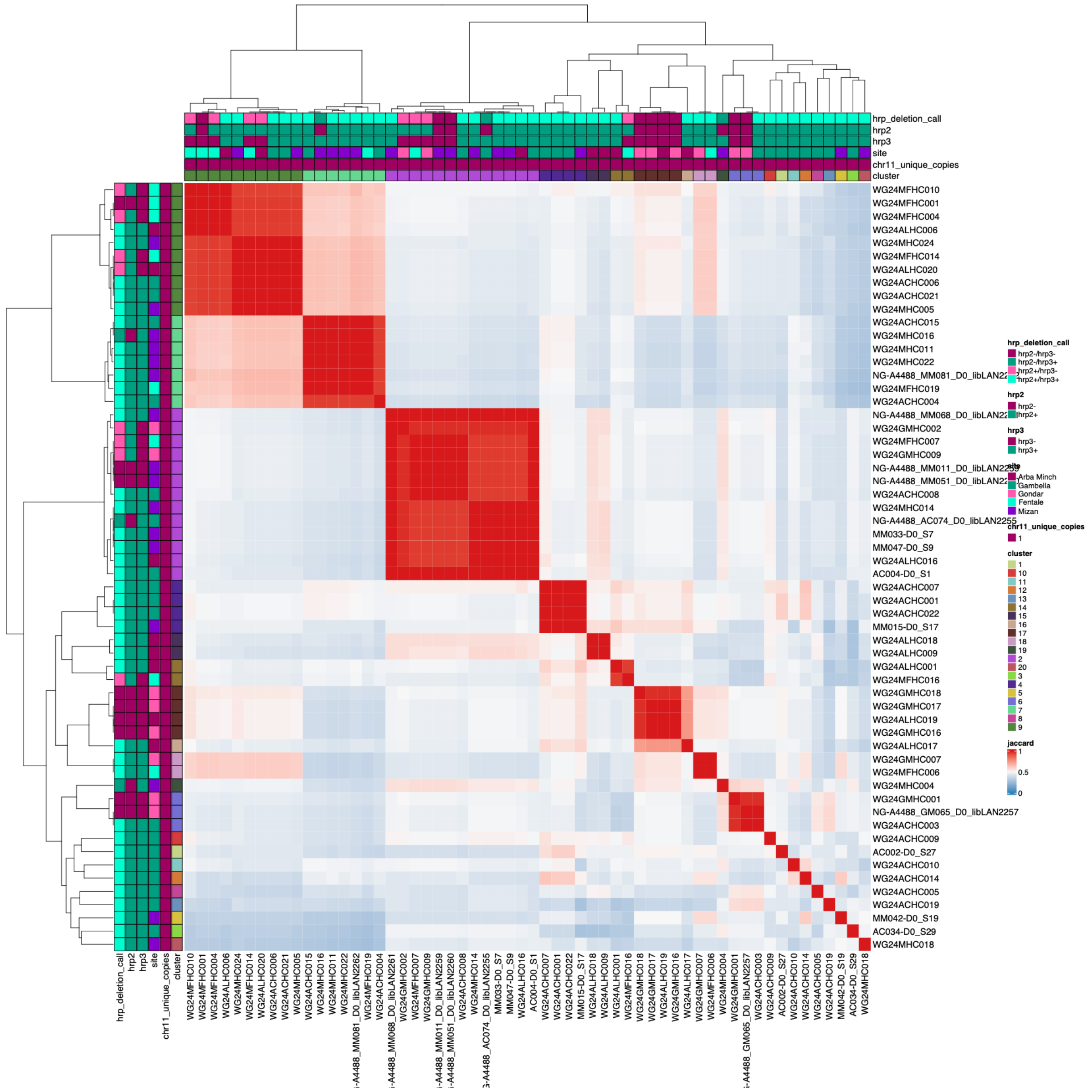

*Pairwise haplotype sharing across chromosome 11 targets, expressed as the Jaccard index, among P. falciparum monoclonal infections harbouring a single unique chr11 haplotype. Heatmap colour represents the Jaccard index, ranging from 0 (blue; no shared haplotypes) through 0.5 (white) to 1 (red; identical chr11 haplotype repertoires). Rows and columns are ordered by hierarchical clustering of pairwise Jaccard distances (1 − Jaccard), shown as dendrograms. Coloured annotation tracks along the top and left margins indicate, for each sample: combined hrp2/hrp3 deletion genotype, hrp2 deletion status alone, hrp3 deletion status alone, collection site, chr11 unique copy number, and cluster membership.*

**Fig S23. Jaccard index between isolates with 2 unique copies of chromosome 11**

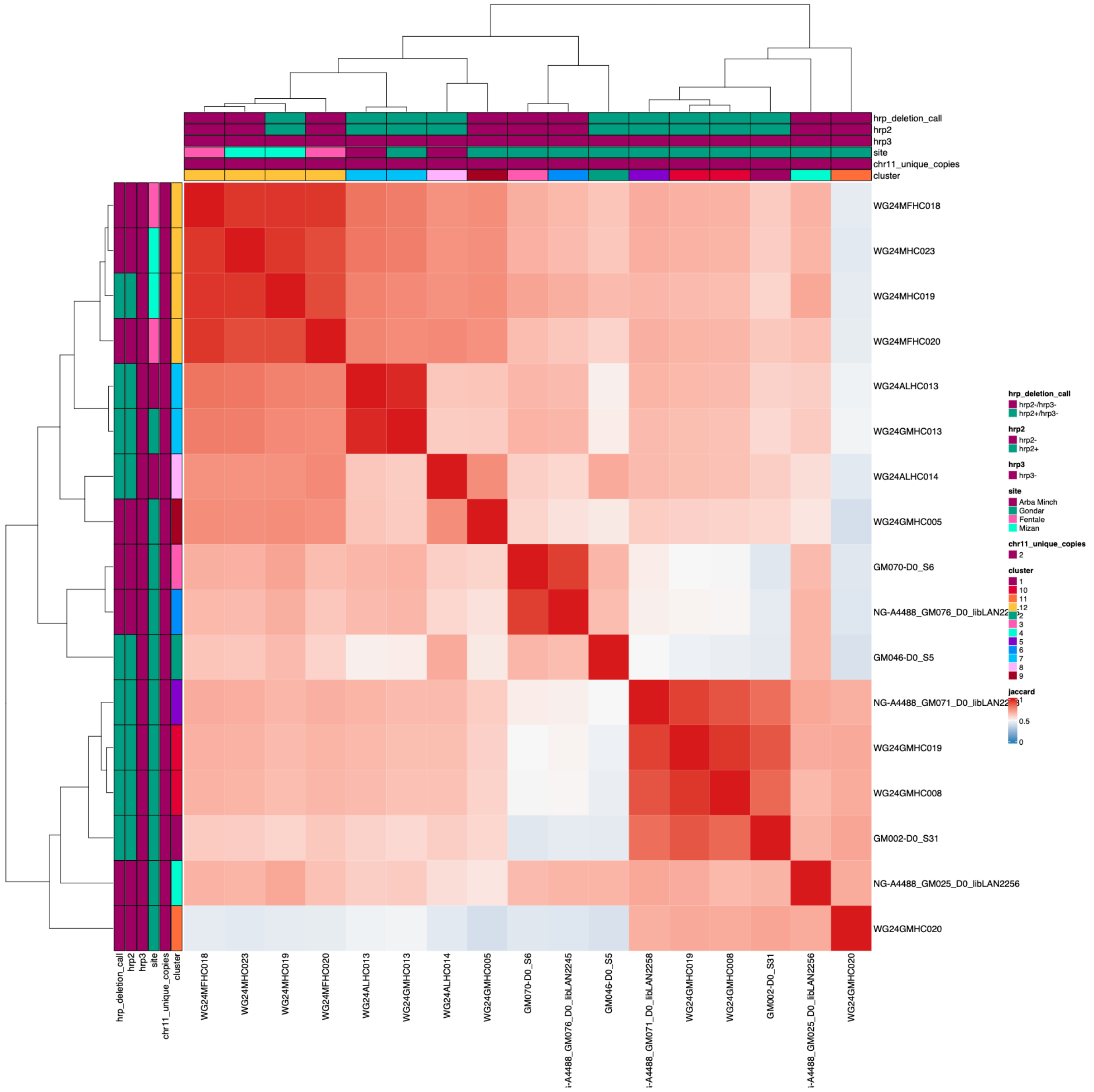

*Pairwise haplotype sharing across chromosome 11 targets, expressed as the Jaccard index, among P. falciparum monoclonal infections harbouring two unique chr11 haplotypes. Heatmap colour represents the Jaccard index, ranging from 0 (blue; no shared haplotypes) through 0.5 (white) to 1 (red; identical chr11 haplotype repertoires). Rows and columns are ordered by hierarchical clustering of pairwise Jaccard distances (1 − Jaccard), shown as dendrograms. Coloured annotation tracks along the top and left margins indicate, for each sample: combined hrp2/hrp3 deletion genotype, hrp2 deletion status alone, hrp3 deletion status alone, collection site, chr11 unique copy number, and cluster membership.*

### Fig S24. Investigation of EHH decay around *Pfhrp2/3* deletions

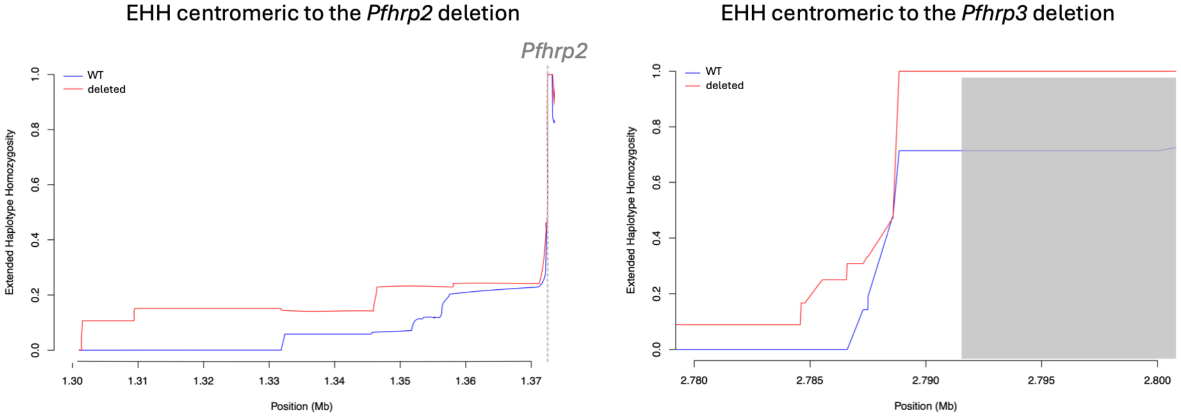

*Extended Haplotype Homozygosity centromeric to the Pfhrp2 and Pfhrp3 deletions in the 2024 Ethiopian isolates. Vertical dashed line indicates the centromeric end of the Pfhrp2 deletion. Grey box indicates the duplication region on chromosome 13. WT and deleted alleles are defined separately for each gene: ancestral denotes an intact copy of the gene in question, derived denotes its deletion. A sample classified as ancestral for one gene may still carry a deletion of the other.*

**Fig S25. Pairwise IBD fractions in wild-type and HRP deleted isolates, per district**

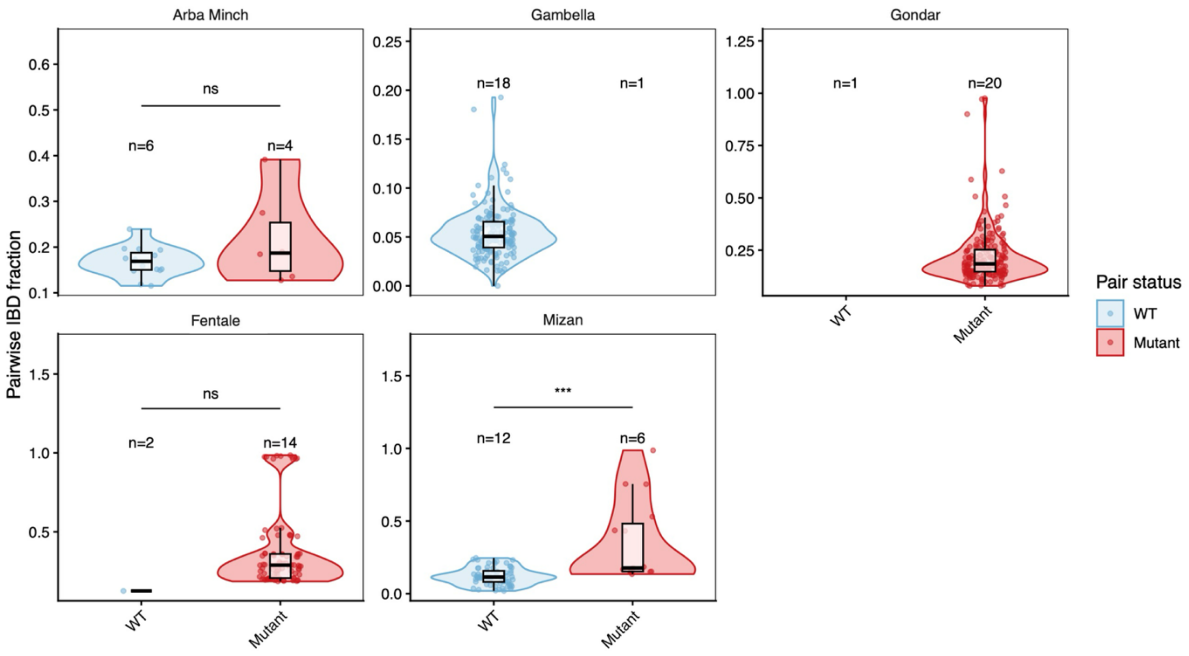

*Violin plots show the distribution of pairwise IBD (fraction of the genome shared IBD) between parasite monoclonal samples within the same district. Comparisons are restricted to pairs where both samples are from the same district and have known Pfhrp2/Pfhrp3 status. Pairs are grouped as wild type (WT; both parasites Pfhrp2+/Pfhrp3+) or mutant (both parasites carrying at least one deletion). Mixed pairs (one wild type and one mutant) are excluded. Overlaid boxplots indicate the median and interquartile range, and individual points represent pairwise comparisons. Numbers above each group indicate the number of individual samples contributing to each category per district. Facets correspond to districts, with y-axis scales allowed to vary between districts. Wilcoxon rank-sum test (*p < 0.05, **p < 0.01, ***p < 0.001, ns = not significant).*

**Fig S26. Multi-tiered analytical framework for the detection of positive selection**

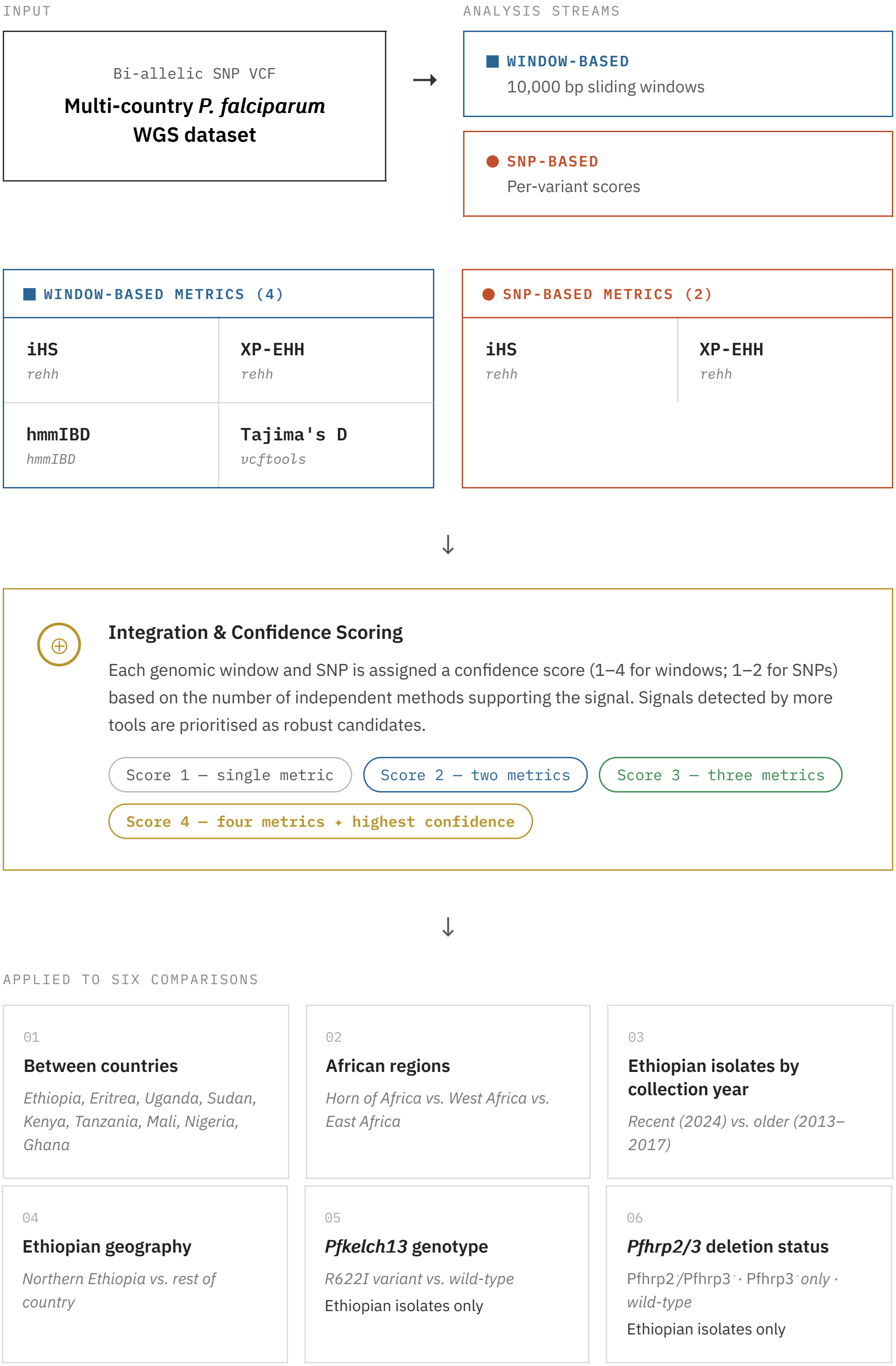

*This flowchart was generated using Claude Sonnet 4.6 from text-based prompt and subsequently edited and verified by the authors. The flow describes the multi-metric selection analysis pipeline applied to a multi-country P. falciparum whole-genome sequencing dataset.*

**Fig S27. Data pruning prior to selection analyses**

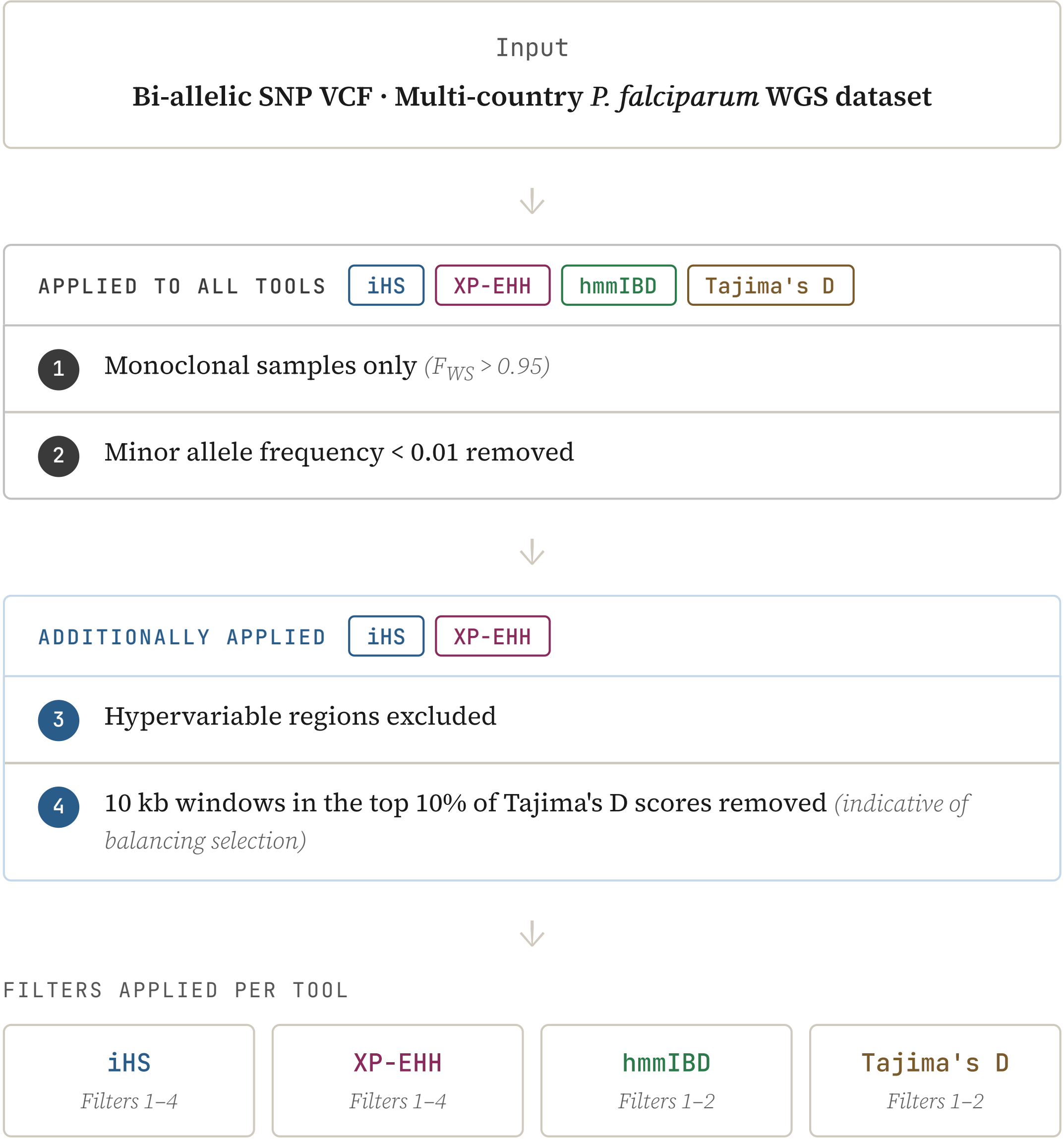

*This flowchart was generated using Claude Sonnet 4.6 from text-based prompt and subsequently edited and verified by the authors. The flow describes the data filtering steps prior to selective pressure analyses.*

### Table S1. Samples included in analysis

| Country | Collection year | Sites | n | Source |
|---|---|---|---|---|
| Ethiopia | 2013 | West Arsi Zone | 11 | [1] |
| | 2015 | Shewa Robit Town | 10 | |
| | 2017 | Atseged Tsimbila, K/Humera, West Armachiho | 17 | [2] |
| | 2017/2018 | | 1 | |
| | 2018 | Atseged Tsimbila, K/Humera | 2 | |
| | 2024 | Arba Minch, Gambella, Gondar, Fentale, Mizan | 120 | this study |
| Eritrea | 2016 | Akordat, Guluj, Shambuko, Tokombia | 47 | [3] |
| | 2017 | Guluj, Shambuko, Tokombia | 26 | |
| | 2019 | Akordat, Guluj, Shambuko, Tokombia | 102 | |
| Sudan | 2015 | Kassala | 5 | [1] |
| | 2016 | Khartoum | 5 | |
| | 2017 | Khartoum | 57 | |
| Tanzania | 2014 | Kigoma, Tanga | 211 | [1] |
| Kenya | 2012 | Kilifi | 60 | [1] |
| | 2013 | | 1 | |
| | 2014 | Kilifi, Kisumu | 68 | |
| | 2015 | Mfangano, Suba | 29 | [4] |
| | 2018 | Bungoma, Chwele, Kibuke, Kilifi, Kimilili, Lukhuna, Luucho, Makhonge, Malindi, Matibo, Mukhweya, Mupeli, Musese, Nabukhisa, Sanandiki, Sikulu, Sikusi, Sirisia, Teso | 31 | [1,5] |
| | | Ngodhe | 10 | [5] |
| Uganda | 2010 | Apac | 12 | [1] |
| | 2017 | Agago, Lamwo | 8 | [6] |
| | 2018 | Busia | 4 | |
| | 2019 | Agago, Busia, Kaabong, Katakwi, Lamwo, Tororo | 20 | |
| | 2020 | Agago, Kaabong, Katakwi, Kole, Lamwo | 65 | |
| | 2021 | Agago, Mbale, Tororo | 47 | |
| | 2022 | Agago, Mbale | 7 | |
| Ghana | 2017 | Greater Accra, Upper East | 40 | [1] |
| | 2018 | Eastern, Upper East | 60 | |
| Nigeria | 2017 | Lagos | 66 | [1] |
| Mali | 2017 | Koulikoro, Sikasso | 14 | [1] |
| | 2019 | Morodjambougou, Moussadjiebougou, N'Korobougou, N'tentoukoro, Soussounkoro, Tenemambougou, Zelani | 42 | [7] |
| | 2020 | Djalakoro-Djitoumou, M'Pana, Morodjambougou, N'Korobougou, N'Tentoukoro, Sikoro-Djitoumou, Tamala, Tenemambougou, Zelani | 22 | |
| India* | 2017 | Odisha, Tripura, West Bengal | 84 | [1] |
| | 2018 | Odisha, West Bengal | 14 | |
| Myanmar** | 2011 | Tanintharyi | 2 | |
| Thailand** | 2008 | Tak | 6 | |
| | 2011 | Tak | 5 | |
| | 2012 | Tak | 6 | |
| | 2013 | Tak | 3 | |

*Samples included in analyses conducted in this paper. *Indian isolates were only included in population genetics analyses, including PCA and ancestry. **Isolates from Myanmar and Thailand were selected for having the PfK13 A675V variant and were exclusively used for comparison of flanking haplotypes around the A675V variant. Metadata per sample can be found in **Supplementary Data 1***
